# Enhancing high-temperature degradation of polyethylene terephthalate through a synergistic division of enzyme labour between a solid-degrading thermostable cutinase and a reaction intermediate-degrading thermostable carboxylesterase

**DOI:** 10.1101/2022.02.02.478778

**Authors:** Arpita Mrigwani, Bhishem Thakur, Purnananda Guptasarma

## Abstract

Cutinases degrade PET (polyethylene terephthalate) into various degradation intermediates (DIs) such as OET (oligoethylene terephthalate), BHET (bis-hydroxyethyl terephthalate), and MHET (mono-hydroxyethyl terephthalate), and eventually into TPA (terephthalic acid), which is the terminal product of degradation. Unlike PET, which is insoluble, TPA and the DIs are sparingly soluble in water. This causes both DIs and TPA to be partitioned into aqueous solution, where DIs accumulate without undergoing significant further degradation, despite being better substrates of cutinase than solid PET. This frustrates the creation of a circular economy involving PET and TPA (since TPA must be separated from DIs before re-condensation into PET). We argue that the non-degradation of DIs owes to cutinase becoming progressively depleted from solution, through binding to solid PET. This creates a conundrum, in that degradation of PET is anticipated to be inversely correlated with degradation of DIs (at least while solid PET remains available to deplete cutinase from solution), causing any improvement of the cutinase’s PET-binding efficiency to only further ensure non-degradation of released DIs. Here, we propose the deployment of a second DI-degrading enzyme; one that remains in solution, and acts as an ‘assistant’ to the ‘master’ PET-invading cutinase acting upon PET’s surface. We demonstrate that one such dual-enzyme system, consisting of a thermostable *Thermus thermophilus* carboxylesterase (TTCE), characterized here for the first time, and the already-used thermostable leaf-branch compost cutinase (LCC), allows complete degradation of all products of PET hydrolysis into TPA in solution, at 60 °C, even in the presence of residual solid PET.

## Introduction

PET (polyethylene terephthalate) is a plastic and an environmental pollutant that is currently accumulating around the world, at alarming rates. The need exists to urgently develop methods for the degradation of PET at rates comparable to those at which it is being generated. One of the modes of degradation that is seriously being explored involves the use of enzymes. As PET is a human creation, microorganisms have not yet evolved to efficiently use it as a carbon source by degrading it enzymatically. Even so, some microorganisms like *Ideonella sakaiensis* appear to have evolved to deploy existing enzymes (specifically, carboxylesterases (CEs) or lipases) to degrade PET and utilize it as a carbon source,^1^ by effectively exploiting the fact that PET’s ester bond-containing backbone is vulnerable to degradation by any enzyme that is capable of hydrolysing ester bonds. Therefore, current efforts aimed at developing and improving methods for the enzymatic degradation of PET are uniformly focused upon the use and/or improvement of CEs, and other related enzymes, not just from *I. sakaiensis*,^2-5^ but also from other sources.^6-9^

Carboxylesterases can be either general purpose CEs, or CEs with preferences for certain types of substrates, e.g., lipases, or cutinases.^10^ All CEs belong to a superfamily of esterase enzymes that display diverse functions,^11^ synthesizing or trans-esterifying ester bonds, or hydrolysing carboxylic esters, thioesters and amides, to release carboxylic acid and alcohol, as by-products.^12^ CEs are produced by every known form of life, including microbes that grow at high temperatures. When CEs act upon PET, they hydrolyse several different kinds of ester bonds. On the one hand, they can engage in ‘endolytic’ action at sites located in the middle(s) of PET chain(s), to generate short oligo-ethylene terephthalate (OET) molecules containing only a few units of ethylene terephthalate. On the other hand, CEs can also engage in ‘exolytic’ action upon the termini of PET chains, or OET chains, to either release TPA [terephthalic acid; also known as terephthalate] which happens to be the terminal product of all degradation of PET or, in addition to TPA, also certain degradation intermediates (or DIs) such as MHET [mono-hydroxyethyl terephthalate, consisting of an ethylene glycol (EG) linked to TPA], or BHET [bis-2-hydroxyethyl terephthalate, consisting of two ethylene glycol (EG) groups linked to TPA, with one EG placed on either side].^13^ The structures of PET, BHET, MHET, TPA, and EG, are shown in Figure S1 †.

Typically, enzymatic degradation of PET produces both TPA and copious amounts of DIs such as OET, BHET and MHET. Although, to the best of our knowledge, this has not been discussed explicitly in the literature, we begin this paper by stating explicitly that we consider it to be entirely likely that the accumulation of DIs occurs because PET is a solid substance which is hydrophobic and insoluble, whereas all of the DIs of PET are sparingly soluble in aqueous media, in comparison. The higher solubility of the DIs thus causes the partitioning of the DIs away from solid PET, into aqueous solution, concomitantly with their production by enzymes engaged in the degradation of solid PET. The main consideration, in this regard is the following. An enzyme that is required to bind to, and invade, solid PET must itself be likely to become progressively depleted from the solution surrounding solid PET, owing to its titration by the surface of PET, through binding of PET (which is something that is required for the degradation of PET, in any case, since any enzyme that remains in solution, without undergoing binding to PET, cannot ever degrade PET which is presented to it as a solid). The consequence of this consideration is that there is likely to be a severe lack of availability of PET-hydrolysing enzyme in the solution that surrounds any solid PET undergoing degradation, if the only enzyme that is used is a single enzyme that has been perfected to bind to, and degrade, solid PET. In other words, the more perfect the enzyme is at binding to PET, invading it, and hydrolysing it into DIs and TPA, the higher is the likelihood that the enzyme would be unavailable in solution, to effect the degradation, into TPA, of the OET, BHET and MHET that have escaped away from solid PET into the aqueous solution.

We emphasize that it is this unavailability of enzyme (e.g., a very efficient PET-degrading cutinase) in solution to degrade DIs (caused by the gradual and progressive titration of all of the PET-hydrolysing enzyme population onto PET’s surface, during a hydrolytic reaction) which is likely to be primarily responsible for the accumulation of DIs in solution, i.e., DIs that remain largely undegraded, at least for as long as any solid PET remains available. The most important implication of the above argument is that the enzymatic degradation of DIs into TPA can be expected to vary in inverse proportion with the efficiency of enzymatic degradation of solid PET by a single enzyme that has been perfected to bind to, and invade, PET. Although this argument providing a rationale for the accumulation of DIs has not been discussed explicitly in the literature, it is certainly a well-acknowledged fact that the accumulation of DIs is a problem (discussed in detail in Section 1 of the Supporting Information). This is because it is expensive to separate DIs from TPA, and this frustrates the creation of a circular economy involving PET and TPA, based upon the recycling of TPA into fresh PET.^14^ An additional consequence of the non-existence of this circular economy is the higher dependence of PET-manufacturing processes upon petroleum/fossil fuels.

It is widely recognized that an urgent need exists to improve enzymes, and enzyme-based processes, towards achieving the complete (and ‘Green’) degradation of PET into TPA, with no residual DIs. In Section 1 of the Supporting Information, a detailed discussion of all of the advantages of doing so, in respect of every aspect of enzymatic degradation of PET that can be related to ‘Green Chemistry’, together with the advances offered by the work presented here, is provided along with a detailed list of references (distinct from those provided in this manuscript). Thus far, although a dual-enzyme system has been developed with the purpose of reducing MHET-based inhibition of cutinase enzyme through degradation of MHET (discussed in Section 1 of the Supporting Information, and discussed later in this introduction; see reference number 18), no attention has yet been given to deploying an additional enzyme to assist the main enzyme that is deployed to invade PET’s solid surface, to improve both TPA yield and purity, i.e., as ‘assistant’ enzyme which mainly degrades OET, BHET and MHET into TPA, without displaying any significant binding to PET. This is the lacuna that we address here.

We explore the use of two different types of CEs to simultaneously, and synergistically, perform two different types of PET degradation, in two separate environments that happen to be always present around solid PET, in any reaction attempting the enzymatic hydrolysis of PET. These environments are: (i) the solid-liquid interface, at which a ‘master’ enzyme can bind to solid PET, to perform the initial hydrolysis of PET into TPA, and DIs, and (ii) the solution around solid PET, in which an ‘assistant’ enzyme can degrade the dissolved DIs generated by the master enzyme, into TPA. The intended consequences of this engineered division of labour are an improvement in the overall yields of TPA from the degradation reaction (i.e., TPA productivity), and a reduction in the accumulation of DIs (i.e., TPA quality). We demonstrate that improved yields of TPA, as well as almost-pure TPA, can indeed be obtained through the judicious selection of two synergistically-acting enzymes that divide up the labour of degrading PET into two separate types of degradation tasks; one at the surface of solid PET, and the second in the solution around solid PET. As the ‘master’ enzyme, we have selected LCC (leaf branch compost cutinase), a well-known enzyme derived from a metagenomic library which is currently acknowledged to be the most PET-invasive enzyme thus far known, or characterized.^6^ LCC is known to have a temperature of optimal function of ∼70 °C, allowing it to work in the range of temperatures at which most forms of PET display a glass transition temperature.^15^ At temperatures approaching or exceeding this temperature, PET is expected to expose the backbones of individual polymer chains to a greater extent to the surroundings. LCC has a significantly hydrophobic surface which presumably helps it to bind to PET, which is highly hydrophobic, but which also causes LCC to aggregate in aqueous solution, when it is used at moderately high concentrations. Attempts have been made to reduce LCC’s tendency to aggregate, through rational mutagenesis and through glycosylation,^16^ and to improve its thermal stability further through the rational introduction of a disulphide bond.^6^ Random mutagenesis has also been used to improve LCC’s catalytic efficiency through combinatorial site-saturation studies.^6^

As an ‘assistant’ enzyme, we have selected a previously-uncharacterized enzyme that we call TTCE (*Thermus thermophilus* carboxylesterase). TTCE (or TT1662) is derived from the proteome of strain HB8 of *T. thermophilus*. TTCE’s three-dimensional structure (RCSB ID: 1UFO) has been determined and deposited in the Protein Data Bank, as a part of a structural proteomics initiative.^17^ However, it has not been previously characterized in any other way, and the presence of esterase activity in TTCE is being described for the first time in this manuscript. The structure of TTCE has a canonical α/β fold in which a hydrophobic core constituted of eight parallel β-strands is surrounded by six α helices. A presumed triad of TTCE’s catalytic residues has been determined through comparison of its structure with that of a structurally-related enzyme from *Pseudomonas fluorescens*, and this triad is thought to consist of a conserved histidine residue, a conserved (acidic) aspartic acid residue, and a conserved (nucleophilic) serine residue.^17^ We have produced and characterized TTCE (as described in this paper) and found it to be extremely thermally-stable, extremely chemically-stable, non-aggregation-prone, poorly capable of binding to (or invading, or degrading) solid PET, moderately efficient at turning BHET into TPA and EG, in solution, and characterized by a preference for acting upon short chain aliphatic esters and small aromatic esters. This combination of characteristics causes TTCE to almost perfectly fit the description of an ideal ‘assistant’ esterase to a PET-invading, thermostable cutinase like LCC, since TTCE could be expected to largely remain in solution and degrade only DIs into TPA, whereas LCC could be expected to largely remain on the surface of PET and degrade it into DIs, and TPA.

We must emphasize that although this is very likely the first study that uses enzyme synergy to facilitate a clear division of labour (through a partitioning of enzymes between solid and solution) to effect degradation of PET and its DIs, this is not the first time that enzyme synergy has been applied to degradation of PET. In three pioneering studies that have already been conducted,^18-20^ cocktails of different CEs, as well as genetic fusions of different CEs, have been shown to enhance PET degradation. One reason that an approach exploiting enzyme synergy holds promise (over and above reasons already discussed in the previous paragraph, which mainly relate to differences in the relative hydrophobicity/hydrophilicity of surfaces of different enzymes, PET, and the DIs) is also that PET itself contains several different types of ester bonds. These bonds are presented to hydrolytic enzymes in a variety of different three-dimensional configurations, and physical contexts. Certain CEs are efficient at acting upon the termini of PET chains through ‘exo’ polymer-degrading action, whereas most CEs are thought to act upon the middle regions of PET chains, through ‘endo’ polymer-degrading action.^21^ Then again, in respect of both types of action, certain CEs are thought to optimally act upon PET at temperatures below the glass transition temperature of crystalline PET, while other CEs are thought to act upon semi-crystalline PET, at temperatures above the glass transition temperature of crystalline PET, and while yet other CEs are thought to act only upon non-crystalline (amorphous) PET at or above/below the glass transition temperature of amorphous PET (∼67 °C) which is nearly 10 °C lower than that of crystalline PET.^22^ The use of different enzymes in a synergistic fashion, therefore, is also likely to facilitate the degradation of different ester bonds in PET, in different physical contexts and not just in different chemical contexts.

It may be noted that PET, as a substrate, tends to be characterized by significant surface hydrophobicity, owing to the placement of aromatic groups at regular intervals along the PET backbone, and also because of the three-dimensional arrangement of PET chains in semi-crystalline or crystalline arrangements. Thus, in addition to the need for a PET-hydrolysing CE to possess the appropriate catalytic function, a CE that acts upon PET in films (or in the form of post-consumer packaging/containers) is required to possess significant surface hydrophobicity, both in the vicinity of its catalytically-active site and also in other regions of its surface, to enable it to bind to PET, and to function in the hydrophobic environment that PET’s surface presents to the aqueous solution.^23^ CEs that are required to act only upon DIs such as OET, BHET and MHET, or only upon BHET and MHET, are not expected to possess the necessary surface hydrophobicity to invade PET, and may be expected to only possess hydrophobic surfaces in the vicinities of their substrate-binding sites (in order to accommodate the terephthalate moiety in PET-derived DIs) and not in other regions of their surfaces, since such CEs might not be required to bind to solid PET. Due to their lower (overall) surface hydrophobicity, such CEs would also be more likely to be soluble in water, than PET-invading CEs. This is likely to make such CEs capable of reaching much higher enzyme concentrations, and displaying a lower tendency to form any aggregates in solution, than most CEs that are known to be good at invading PET, e.g., LCC, which is aggregation-prone. In other words, different structural and surface characteristics apply to the different types of CEs. It is already known that the dimensions and three-dimensional features of catalytically-active sites in CEs that hydrolyse non-crystalline PET are different from those of CEs that hydrolyse crystalline PET.^24^ Further, it is known that there are differences in the catalytic efficiencies of CEs in carrying out hydrolysis of ester bonds in different substrates,^24^ and that this causes some CEs to be catalytically superior at invading and degrading PET, and others to be efficient at degrading DIs. Clearly, therefore, the case exists to argue that degradation of a substrate like PET requires a variety of enzymes of differing enzymatic capabilities, rather than just the capabilities that might be found in a single enzyme. It appears increasingly unlikely that a single enzyme will ever be gainfully employed to completely degrade PET into TPA. Rather, it seems likely that there will be a need to use a combination of: (i) a robust PET-invading CE, and (ii) a robust ‘assistant’ CE that remains in solution, working to mainly break down DIs released into solution by the robust PET-invading CEs, with minimum direct involvement with PET.

While the bulk of research attention thus far has remained focused on the study and development of single enzymes, rather than dual (or multiple) synergistically acting enzymes, quite remarkably, even lesser attention has been given to the systematic exploration of thermostable or hyper-thermostable enzymes capable of acting at temperatures above PET’s glass-transition temperature(s). These lacunae are simultaneously addressed in the work presented in this paper, through the identification and use of a highly-thermostable CE (TTCE) to degrade DIs produced by a moderately-thermostable CE (LCC; a cutinase). We present data here to demonstrate a useful, as well as significant, synergy of action between TTCE and LCC (in which TTCE acts as an ‘assistant’ to LCC, which acts as a ‘master’ CE). Our data suggests that in a cocktail containing the two enzymes, they complement each other’s functions, with one functioning mainly upon PET’s surface, and the other functioning mainly in solution. Together, a cocktail of TTCE and LCC is significantly better than a single PET-invading enzyme like LCC (∼30-100 % better, depending on enzyme concentrations) at generating TPA from commercially-sourced PET, as well as enormously better than TTCE alone (∼2600 % better). Using enzyme concentrations that are feasibly achieved (e.g., 2 µM LCC and 2 µM TTCE), we show that it is also possible to turn PET into pure TPA, with no detectable DIs. We think, therefore, that such a use of enzyme synergy is the ideal way forward for all green chemistry-based enzymatic degradation of PET, even if the reagents that survive into future research and development are enzymes other than LCC and TTCE.

## Results and discussion

### The binding of LCC and TTCE to small terephthalate-based substrates is predicted to be similar by bioinformatics approaches (docking and MD simulations)

#### Structural similarity of LCC and TTCE

The structures of TTCE and LCC were aligned using the TM-align web server,^25^ and were found to align with a root mean square deviation (RMSD) of only 2.33 Å, for Cα atoms, and a TM-score of 0.711 (a value of ≥ 0.5 being indicative of a high similarity of folds). Structural comparison of the active sites in LCC and TTCE, however, revealed the presence of additional structures in TTCE in the form of an extended α helix, and a loop incorporating a small helix, which lie in the immediate vicinity of the substrate-binding and catalytic sites (just above these sites), as is evident from Figure 1A which presents the alignments of the backbones of the two proteins. In LCC, the said loop/helix structures are absent. This suggests that TTCE has limitations in the sizes of the substrate(s) that it can accommodate in its active site, whereas LCC appears to have an active site that can bind to large substrates. We docked short-chain PET (i.e., OET, or oligo-ethylene terephthalate) consisting of four residues [2HE-(MHET)^4^] of MHET (mono-hydroxyethyl terephthalate) to LCC and TTCE. The similarities of docking scores for the two proteins, obtained using Schrodinger Glide-XP-dock, as well as the similarities in the free-energies of binding of the two proteins to this ligand (−71.95 *vs* -72.93 kcal/mol), obtained using Schrodinger Prime-MMGBSA, suggest that LCC and TTCE bind to 2HE-(MHET)^4^ to similar degrees, with LCC binding only somewhat better than TTCE. Table S1 † summarizes the relevant XP-dock and MMGBSA data, in this regard.

**Figure 1:**
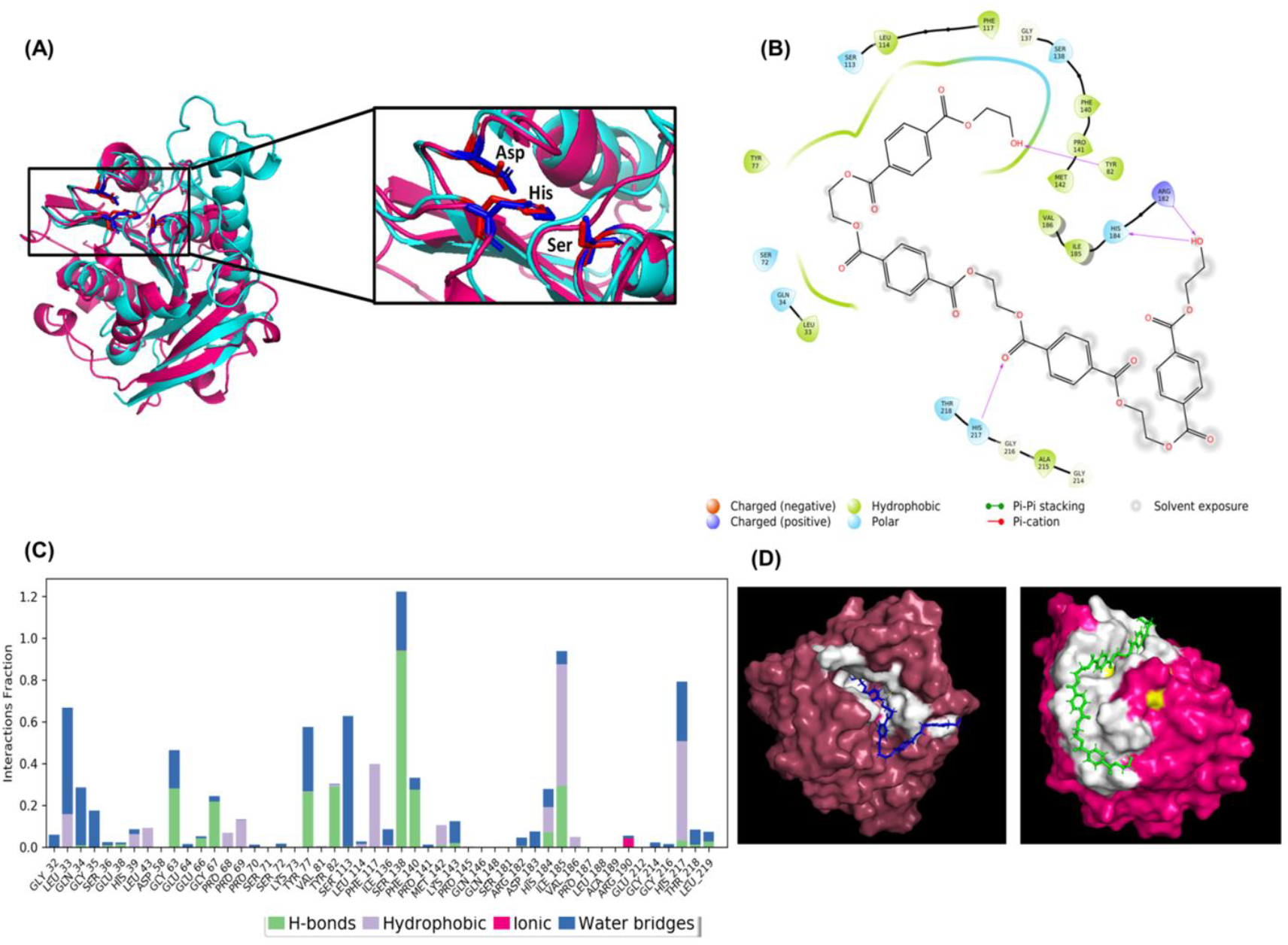
Structural alignment, molecular docking and molecular dynamics simulations. (A) Structural alignment of the backbone of TTCE (cyan, PDB entry 1UFO) with that of LCC (pink, PDB entry 4EB0). (B) Residues surrounding the ligand [2HE-(MHET) _4_] that form the binding pocket of TTCE and interact with the ligand for >30 % of the simulation time. (C) Residues obtained after MD simulation using Simulation Interaction Diagram tool, showing the nature of interaction, i.e., hydrophobic, H-bonding, electrostatic and water bridges as well as the extent of these interactions. (D) Comparison of the 2HE-(MHET) _4-_binding surface of TTCE and LCC. The catalytic residues are shown in yellow and the binding residues are shown in white for both.

Surface representations of 2HE-(MHET)_4_ binding to TTCE and LCC are shown in Figure 1B, while Figures 1C and 1D show the residues that constitute the binding pockets in the two enzymes (as determined from molecular docking and/or MD simulations). Typically, in an MD simulation for a protein-ligand interaction, the stability of the RMSD values after a certain period of simulation time is an indicator of an approach to equilibrium and the compactness of the protein as suggested in Figure S2A† in the supporting information shows that this stability was achieved at a time point beyond 50 ns of simulation with no substantial conformational change in the protein. Figure S2B † plots the fluctuations (RMSF values in Å) suggesting the changes in the local positions of individual residues as a function of simulation time, with β-strands in blue, α-helices in orange, loops in the white. Hydrophobic residues lining the binding pocket in the enzyme(s) appear to participate in π-π stacking interactions with terephthalate groups in the ligand, 2HE-(MHET)_4_, with some water bridges and hydrogen bonds also seen to be formed between the enzymes and the ligand molecules. Figure 1D shows that LCC has a shallower ligand binding surface than TTCE, with the catalytic residue triad and other residues involved in ligand binding clearly accessed with greater ease, by any molecules in the solvent, in LCC, than in TTCE, especially where large substrates are concerned. Further, as already suggested by Figure 1A which shows the structures of LCC and TTCE in the absence of ligand(s), the catalytic residue triad and other residues involved in ligand binding in TTCE can be seen, in Figure 1D, to be located within a deep groove capped by a lid-like structure that is absent in LCC (due to the presence of an additional extended helix, and a loop containing a short helix). From the above analyses, it appears that LCC must be capable of acting upon both long PET chains, as well as upon short PET chains (e.g., OET), due to the shallowness and higher accessibility of its substrate-binding site. In contrast, it appears that TTCE must be capable of acting mainly (or even only) upon OET chains, BHET and MHET, with insignificant action upon long PET chains, due to the fact that its active site lies within a deep groove that is capped, furthermore, by some loop/helix structures.

### The binding of LCC and TTCE to PET is predicted to be different (by structural considerations) and also demonstrated to be different (by binding experiments)

Figures 2A and 2B show the features and characteristics of LCC and TTCE in the region of the predicted active site and surrounding areas, in the absence of any ligand. From these, it is very clear that the hydrophobicity of LCC’s surface is very much higher than that of TTCE, both in the regions immediately adjoining, or constituting, the substrate binding and catalytically-active site, as well as in neighbouring regions of the surface. This leads us to predict that LCC would display much higher binding to a highly hydrophobic solid substance like PET, than TTCE. We examined this possibility experimentally by incubating the two enzymes with PET for a few tens of hours at 60 °C.

**Figure 2:**
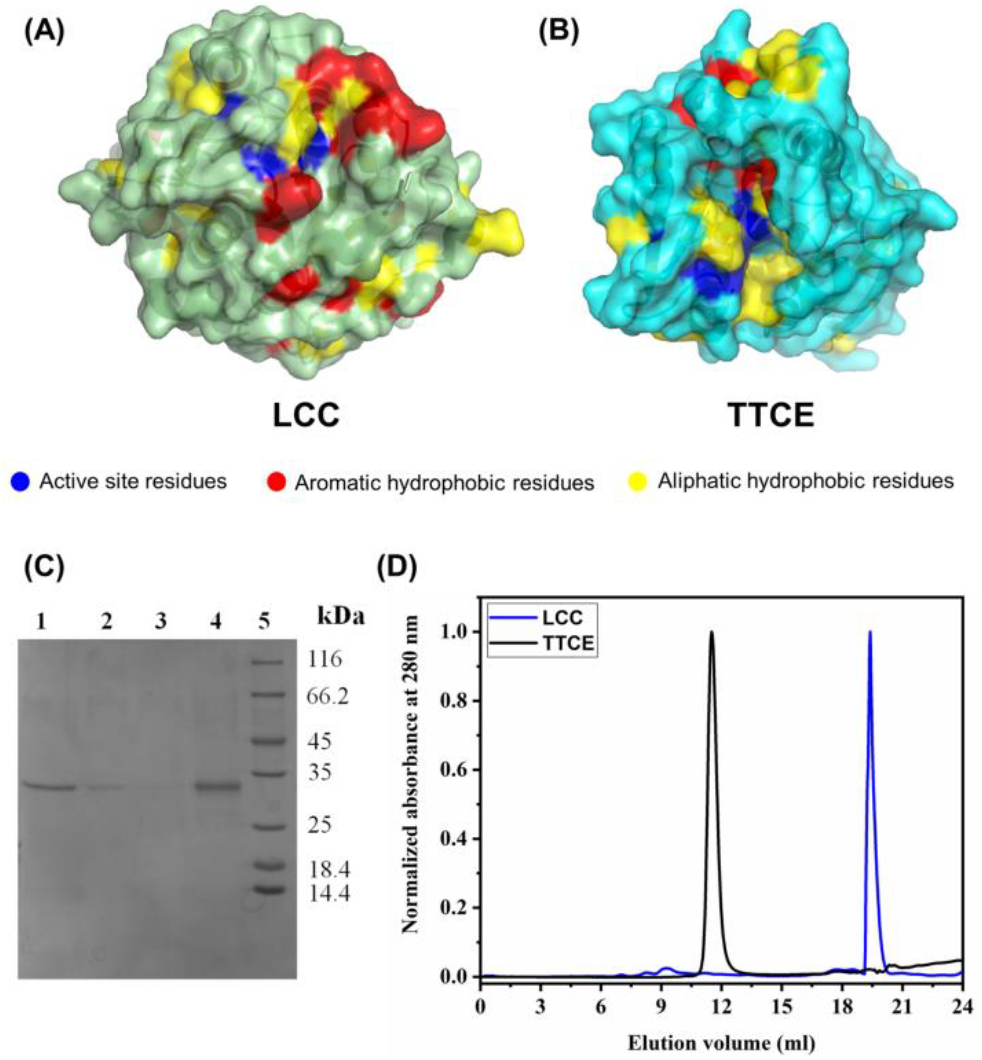
Predicted and observed differences in the PET-binding behaviour of LCC and TTCE. (A) Hydrophobic regions (in red) surrounding the catalytic sites (in blue) on the surface of LCC (in green). (B) Hydrophobic regions (in red) surrounding the catalytic sites (in blue) on the surface of TTCE (in cyan). (C) SDS-PAGE analysis performed to determine PET-binding and partitioning (between PET’s surface and aqueous solution) of TTCE and LCC, after 40 hours of incubation of enzyme with a 6 mm diameter disc of commercially sourced PET film. Lane 1: TTCE left in solution after incubation; Lane 2: TTCE bound to PET’s surface after incubation (extracted through boiling with SDS-PAGE loading buffer, after washing of the film); Lane 3: LCC left in solution after incubation; Lane 4: LCC bound to PET’s surface after incubation (extracted in a similar manner to that shown in Lane 2); Lane 5: Protein molecular weight markers. (D) Elution profiles of TTCE (in black) and LCC (in blue) during gel filtration chromatography on a Superdex-75 Increase (GE) column. The molecular weight of TTCE is ∼26.9 kDa, while that of LCC is ∼29.4 kDa.

We then used SDS-PAGE analyses to assess the differential extents to which the enzymes either (a) remain in solution, or (b) become required to be dissociated from the surface of PET, through boiling in the presence of SDS, for subsequent visualization on SDS-PAGE. Figure 2C shows the results of such an SDS-PAGE (representative) experiment conducted to examine these differential tendencies. After 40 hours of incubation, supernatants and PET films were separately boiled with SDS-PAGE loading buffer (SLB), and samples were loaded onto SDS-PAGE to assess differences in the amounts that remained in solution, or became required to be deliberately dissociated away from the film to which they had potentially undergone binding. Lanes 1 and 2 of Figure 2C show that the bulk of the TTCE enzyme does indeed remain in solution, as predicted, with very little binding occurring to the PET film. In contrast, lanes 3 and 4 of Figure 2C show that the bulk of the LCC enzyme become associated with the PET film, as predicted, with an almost undetectable amount of LCC remaining in solution. Notably, these results indicate a stark and significant difference in the PET-binding characteristics of LCC and TTCE, lending support to the intent outlined in the introductory section of this paper, in which we proposed a need to use one enzyme that binds effectively to PET, to invade it, and a second (assistant) enzyme that remains in solution, even in the vicinity of PET, without undergoing almost any binding to PET. The SDS-PAGE experiments make it clear that LCC partitions onto the surface of the solid PET substrate by adsorbing onto the surface of PET (owing to the greater amount of surface hydrophobicity) and offering higher scope for invasive localized degradation of PET by LCC (ostensibly due to LCC’s shallower active site, and lack of restrictions on sizes of substrates). SEM images are presented to support this, in a later section of the paper. At the same time, the SDS-PAGE experiments also make it clear that TTCE remains in solution (owing to its lack of extensive surface hydrophobicity), offering higher scope for disseminated degradation of short PET chains, BHET and MHET (ostensibly due to TTCE’s deeper active site, and lid-like structures covering the site).

Figure 2D presents further evidence for the presence of extensive surface hydrophobicity in LCC, through the use of gel filtration chromatography resins/media. TTCE is observed to elute at an elution volume of 11.7 ml, which is the volume at which it is expected to elute (as a population of a ∼26.9 kDa polypeptide existing in a monomer-dimer equilibrium; as demonstrated in the following section). In contrast, LCC (∼29.4 kDa), which has a similar monomeric size to TTCE (∼26.9 kDa), and which is established to migrate on SDS-PAGE to a similar extent to TTCE, as already demonstrated by Figure 2C, however, turns out to elute at the very much higher elution volume of 19.5 ml. Theoretically, such an elution volume is only possible for LCC if it were to interact with the Superdex-75 Increase (GE) gel filtration chromatographic resin, so as to engage in a combination of gel filtration chromatography (intended) and non-specific affinity chromatography (unintended), causing its elution volume to become significantly displaced, as well as unrepresentative of polypeptide subunit size. Notably, proteins that interact with gel filtration chromatographic resins (e.g., Superdex resins), elute at late elution volumes (and sometimes even at elution volumes exceeding the column’s bed volume), with such proteins typically having very high surface hydrophobicity.

### TTCE is a well-folded enzyme that is highly stable to heat and denaturants (making it an ideal assistant to LCC, at temperatures approaching PET’s glass-transition temperature)

The structure of TTCE is known, due to its having been subjected to crystallization in a structural proteomics initiative during the early years of this century. The function of TTCE is also anticipated to be that of a carboxylesterase, due to the homology of its sequence with that of a carboxylesterase from *Pseudomonas fluorescens*. However, TTCE has remained an uncharacterized enzyme, in which esterase activity has remained to be either demonstrated, or characterized. Furthermore, even though the sequence of TTCE is derived from the proteome of a thermophile microorganism, *Thermus thermophilus*, it is not yet known whether the enzyme is capable of folding into a thermostable structure possessing enzymatic activity, upon heterologous production in *Escherichia coli*. Further, if TTCE were to indeed turn out to be thermostable, the extent of its stability has remained unknown. Therefore, we proceeded to first confirm, and characterize, TTCE’s identity, its thermostability, its thermal stability, its thermodynamic stability, its chemical stability, its kinetic stability to denaturation, and also its activity, as an esterase, against small aliphatic and aromatic substrates, including terephthalate-based substrates.

#### The recombinant ∼26 kDa protein is TTCE

Purified TTCE was electrophoresed on SDS-PAGE, and the band displaying mobility corresponding to a mass of ∼26 kDa was excised and subjected to mass spectrometric analysis, as shown in Figure S3A †. The masses of peptide peaks obtained experimentally were matched with the masses of peptide peaks generated through *in silico* tryptic digestion of TTCE. In Figure S3B †, peaks marked with red arrows identify the many masses that matched the expected (*in silico* digestion-generated) masses, to an accuracy of 1 Da, thus confirming TTCE’s identity.

#### TTCE is predominantly dimeric

We assessed the quaternary structural status of TTCE through a combination of size exclusion chromatography (SEC), glutaraldehyde crosslinking, and dynamic light scattering (DLS) experiments. The chromatogram in Figure S3 C † shows that TTCE elutes at 11.51 ml from a Superdex-75 Increase column, which places its native molecular weight between ∼26 kDa and ∼52 kDa, i.e., between the elution volumes expected for a monomer and a dimer, towards the dimer. That TTCE exists as an equilibrium population of monomers and dimers was also established through crosslinking by glutaraldehyde (a homo-bi-functional chemical reagent that covalently crosslinks proteins placed in close proximity, through reaction of the reagent’s aldehyde group with protein N-termini or the ε-amino groups of lysine residues), as shown in Figure S3D †. Notably, TTCE’s crystal structure provides scope for dimerization through the formation of anti-parallel beta sheets between monomers (13). The hydrodynamic radius obtained by DLS experiments also supports an equilibrium of monomers and dimers. Weight-fractions of population, plotted in Figure S3E † (with the correlation function shown in Figure S3F †), suggest a radius of 3.10 nm (or a diameter of ∼6.2 nm) for 93.51% of the TTCE population, a value which is larger than the radius of a monomer (∼2.4 nm, as reckoned from the crystal structure), and smaller than that of a dimer. Notably, the DLS experiments also showed that a small fraction of the population exists as oligomers or aggregates.

#### TTCE contains a mix of alpha and beta structures

The circular dichroism (CD) spectrum of TTCE is shown in Supporting Information Figure S3G †. It is dominated by α-helices; with low negative intensity at ∼218 nm, owing to the presence of β-sheet content, and high negative intensity at ∼208 nm, owing to helical content and the presence of random coils, which is in agreement with the determined structure of the enzyme.

#### TTCE is extraordinarily thermally stable

We assessed the thermal stability of TTCE through a combination of circular dichroism (CD), fluorescence spectroscopic and differential scanning calorimetric (DSC) studies. Figure 3A shows that there are no variations in the far-UV CD spectrum of purified TTCE as a function of increasing temperature, during heating of the enzyme between 20 °C and 90 °C. The Boltzmann fit of the MRE values at 222 nm plotted against the corresponding temperature indicated that more than 90% of the secondary structure remains intact at 90 °C (data not shown). The monitoring of intrinsic fluorescence derived from TTCE’s single tryptophan, which is expected to display a solvent-dependent, red-shifting of the fluorescence emission maximum value, from the its native value of ∼333 nm (for the folded, and native, protein) to longer wavelengths, owing to protein unfolding, shows that there is no such red shifting observed, as shown in Figure S4A †. This establishes that there are no significant tertiary structural changes accompanying the heating of TTCE, over the same temperature range (from 20 °C to 90 °C). Notably, a gradual drop in fluorescence intensity with temperature is observed, due to the increased probability of thermal de-excitation of the excited tryptophan moiety. The overall conclusion from the above data is that there is effectively no unfolding of the secondary or tertiary structures of TTCE, even at very high temperatures. This conclusion was further tested through micro-calorimetric (DSC) studies. As shown in Figure 3B, the T_m_ from the up-scan (i.e. heating of TTCE from 20 °C to 90 °C) was estimated to be 80.5 °C, with a significant change in enthalpy (8.016*10^6^ Joules/mole) seemingly associated with this structure-melting or unfolding transition, which is suggestive of significant thermodynamic stability in TTCE’s native three-dimensional structure. Notably, the enthalpic transition in DSC is seen at 80.5 °C, although changes in secondary and/or tertiary structure are not noted to occur even up to a temperature of 90 °C, as already mentioned. This suggests that TTCE could exist in a molten globular state at temperatures between 80.5 °C and 90 °C, with the enthalpic transition not being paralleled by any significant change in structure. At any rate, these studies show that TTCE can work at temperatures exceeding the glass-transition temperatures of almost all forms of semi-crystalline and crystalline post-consumer PET (which have been mentioned in the introductory section).

**Figure 3:**
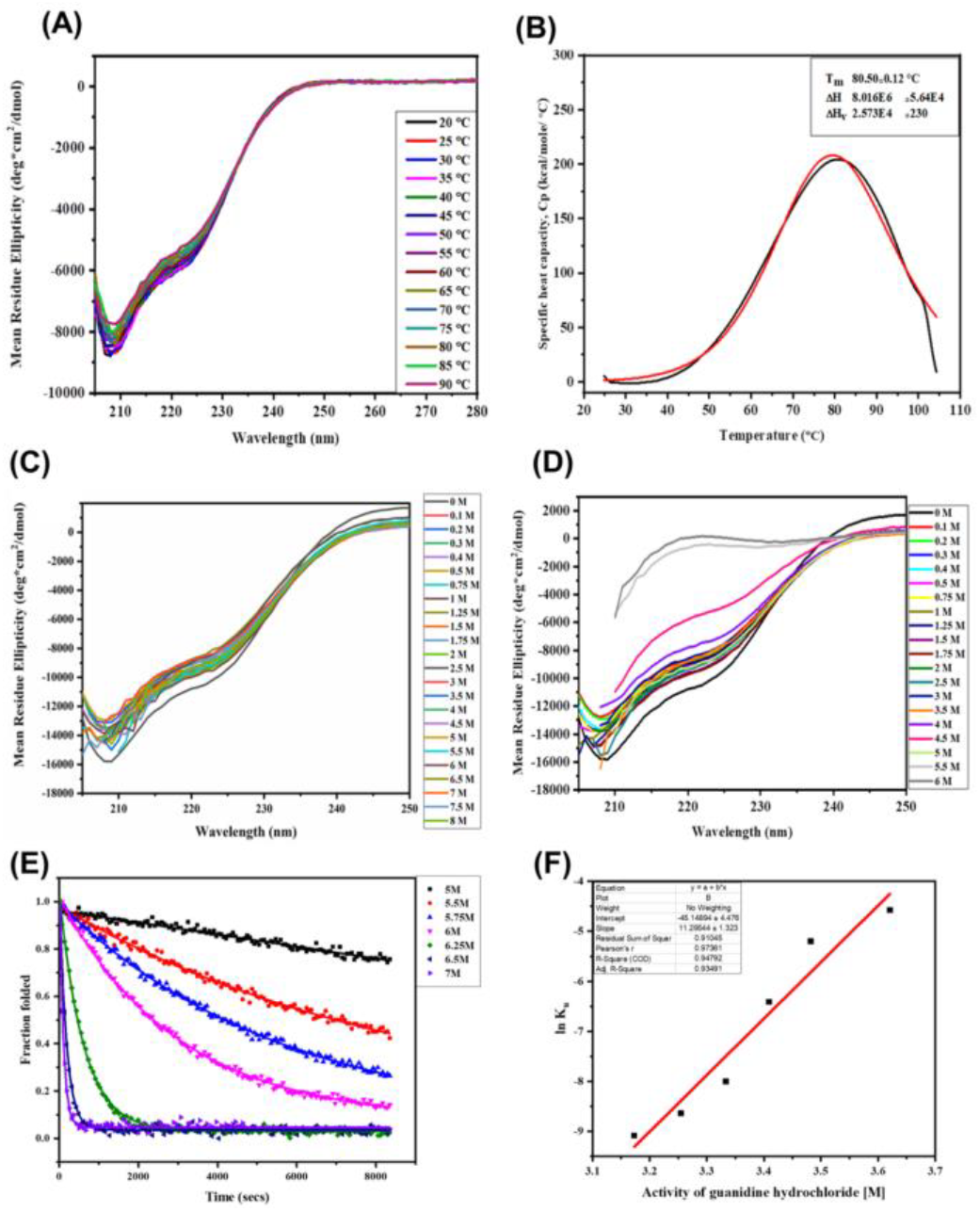
Thermal, thermodynamic, chemical (equilibrium and kinetic) stabilities of TTCE. (A) Lack of changes in secondary structure of TTCE (CD spectra) as a function of heating to different temperatures between 20 °C and 90 °C. (B) Enthalpic changes associated with heating of TTCE (Differential Scanning Calorimetry profile, showing the data in black, and the fitting in red). CD spectra of TTCE incubated overnight with varying concentrations (C) (0 M to 8 M) of urea. (D) (0 M to 6 M) of Gdm.HCl. (E), (F) Kinetic stability studies of TTCE in the presence of Gdm.HCl (also known as Gdm.Cl). (E) Fraction folded versus time plot at Gdm.HCl concentrations in the range of 5 M to 7 M, during the course of 2 hours of incubation with the denaturant. (F) Half-chevron plot obtained from the rates of unfolding determined for different Gdm.HCl concentrations.

#### TTCE appears to be extraordinarily chemically stable, in equilibrium measurements

We next assessed the chemical stability of TTCE through CD and fluorescence studies to explore the energetics of the enzyme’s unfolding at 25 °C, during overnight incubation, by denaturants such as urea, and guanidium hydrocholide (Gdm.HCl). CD data plotting changes in mean residue ellipticity (MRE) as a function of denaturant concentration are shown in Figure 3C and Figure 3D. With urea, Figure 3C and Figure S4B †, respectively, show CD spectra and variations in the intensity of the CD signal at 222 nm, and TTCE is seen to retain the same MRE value at 222 nm over denaturant concentrations ranging from 0.0-8.0 M urea. This demonstrates that even overnight incubation in 8 M urea has no effect upon TTCE’s structure. Figure 3D and Figure S4C † show CD spectra and variations in the intensity of the CD signal at 222 nm with varying concentration of Gdm.HCl. TTCE is seen to retain the same MRE values at 222 nm over denaturant concentrations ranging from 0.0-4.5 M Gdm.HCl, demonstrating that even overnight incubation in 4.5 M Gdm.HCl has no effect upon TTCE’s structure. However, it is observed that 6 M Gdm.HCl completely destroys TTCE’s secondary structure, through a cooperative melting transition with a C_m_ of around 5.1 M Gdm.HCl, as can be seen in Figure S4 C †. Notably, these results are in keeping with the known facts that (a) urea disrupts mainly hydrogen bonds, while Gdm.HCl disrupts hydrogen bonds as well as electrostatic interactions, and that (b) proteins commonly display a C_m_ with urea that is about twice the C_m_ obtained with Gdm.HCl.^26^ The high C_m_ of 5.1 with Gdm.HCl indicates that the C_m_ with urea would be 10.2 M (exceeding the solubility of urea). The data thus indicates that hydrogen bonds determine TTCE’s stability to a lower degree than electrostatic interactions, and also that urea is unable to unfold the protein by attempting to disrupt hydrogen bonds alone. Further, we also examined spectra for intrinsic fluorescence emission owing to tryptophan residues. These are shown as Figure S4D † and Figure S4E †, respectively, for urea and Gdm.HCl. From these panels, it is evident that the tertiary structure of TTCE is largely intact up to a urea concentation of nearly 8 M, and a Gdm.HCl concentration of nearly 4.5 M, indicating that it is not merely TTCE’s secondary structure which is highly stable (as seen in the CD spectra in Figure 3C and Figure 3D), but also its tertiary structure. Notably, the unfolding of TTCE by concentrations of Gdm.HCl exceeding 4.5 M is observed to be accompanied by a dramatic increase in the protein’s fluorescence quantum yield, together with a red shift in the protein’s wavelength of maximal emission, as can be seen in Figure 3D. The increase in quantum yield appears to owe to a change in the environment of tryptophan, which appears to have been released from quenching. The appearance of an emission band at 307 nm appears to owe to cessation of energy transfer between a nearby tyrosine residue (separated by 8 Å from the tryptophan residue), and tryptophan, in the native structure, allowing visibility of the tyrosine’s fluorescence (no longer subject to energy transfer) upon TTCE’s unfolding (see supporting information Figure S4 F †).

#### TTCE also appears to be extraordinarily chemically-stable, in kinetic measurements

To determine the energetic features of TTCE, kinetic studies based on Gdm.HCl unfolding were performed using CD spectroscopy. MRE values at 222 nm, plotted as a function of time (at a particular denaturant concentration, over a period of 2 hours), were used to derive rates of unfolding at each Gdm.HCl concentration, based on the spectra shown in Figures S5A-to-S5G †, covering Gdm.HCl concentrations ranging from 5.0-7.0 M. Subsequently, rates of unfolding of TTCE in the absence of denaturant were calculated from the plot of fraction folded *versus* time, shown in Figure 3E. The slope of the half-Chevron plot (m_u_), presented in Figure 3F, was 11.29. The rate of unfolding of TTCE in water (K_u;w_), calculated from equation (1), was 2.6 ×10^−20^ s^-1^. For comparison, it may be mentioned that many proteins from the hyperthermophilic archaeon, *Pyrococcus furiosus*, have rate constants for unfolding in water (without denaturant) of the order 10^−15^ s^-1^.^27, 28^ The rate of unfolding obtained for TTCE was thus about 5 orders of magnitude slower than that of the proteins from *P. furiosus*, i.e., 2.6×10^−20^ s^-1^. This slow rate is indicative of the extraordinarily high kinetic stability of TTCE to chemical denaturation.

### TTCE acts efficiently upon BHET, but not upon solid PET

#### TTCE acts upon small aliphatic esters and BHET

Figure 4A shows TTCE’s action upon 1-Napthyl butyrate (7.4 µM enzyme; 2 mM substrate), with an optimum temperature of ∼50 °C, with the enzyme retaining 85 % of its maximal activity at 80 °C, owing to its high thermal stability (demonstrated in the previous section). Figure 4B shows TTCE’s action upon an aliphatic long-chain ester, para-nitrophenyl palmitate (1 µM enzyme; 500 µM substrate), with a similar optimum temperature of ∼ 50 °C, with 72 % of its maximal activity retained even at 100 °C. These experiments demonstrate that, upon production in *Escherichia coli*, the amino acid sequence of TTCE folds into a three-dimensional structure that is not merely thermally-stable, but also optimally active at high temperatures as well as active over a wide range of temperatures (including extremely high temperatures), as would be expected for any enzyme designed by nature to fold within *Thermus thermophilus*; of course, only as long as the enzyme were itself able to fold to the designated native structure in a different environment (such as the cytoplasm of *E. coli*) through, e.g., co-translational folding of its polypeptide chain. Many carboxylesterases are known to additionally hydrolyse complex aromatic substrates, and not just small aliphatic substrates. Thus, we decided to test TTCE’s action upon a polyester like PET, which is constituted of repeating units of TPA (terephthalate) and EG (ethylene glycol). We also decided to test TTCE’s action upon BHET, which is a degradation intermediate (DI) of PET. Figure 4C demonstrates TTCE’s ability to act upon BHET (1 µM enzyme; 250 µM substrate; 12 hours; ∼60 °C), in a reaction stopped by 1:1 dilution with acetonitrile, followed by storage in ice prior to HPLC analysis. It was found that under the conditions mentioned (under parentheses) above, TTCE was able to convert ∼97.1 % of the BHET substrate into ∼61.9 % MHET and ∼33 % TPA, with only ∼2.9 % of the original BHET remaining left as residue. Notably, at this temperature, BHET also showed some auto-hydrolysis, generating ∼18.7 % MHET, but no TPA, with ∼81.3 % BHET remaining left as residue. This last result indicates that auto-hydrolysis of BHET is not responsible for the quantitative conversion of BHET into MHET and TPA, by TTCE, and demonstrates the robust action of TTCE upon BHET.

**Figure 4:**
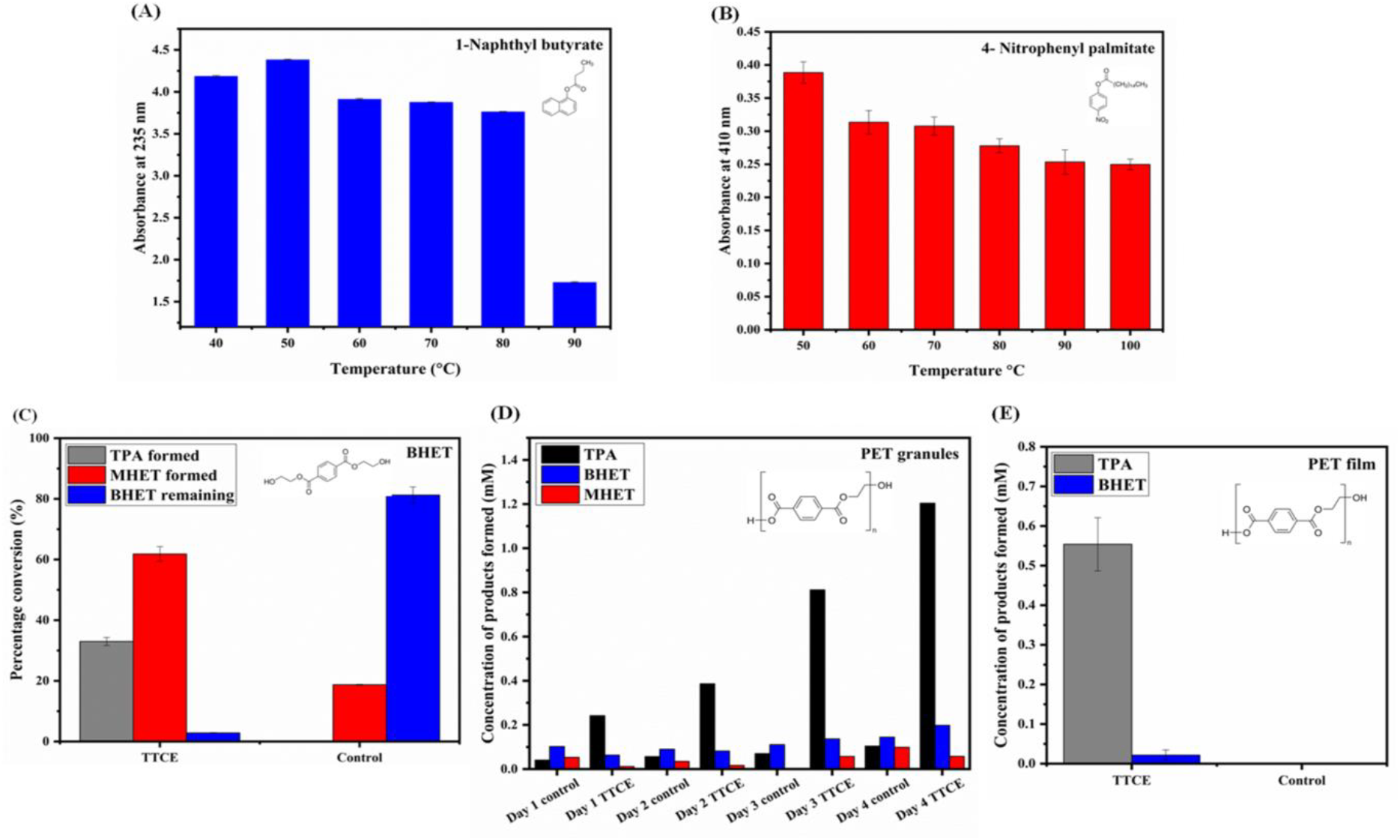
Activity of TTCE upon different substrates. (A) 1-Naphthyl butyrate, quantified through measurement of absorbance at 235 nm. (B) 4-Nitrophenyl palmitate, quantified through measurement of absorbance at 410 nm. (C) BHET (Bis(2-Hydroxyethyl) terephthalate). (D) PET granules. (E) PET film. For C, D, and E, the soluble substrates remaining and the product formed were quantified through reverse-phase HPLC, through monitoring of elution at 240 nm, and measurement of the area of the elution for TPA, BHET and MHET.

#### TTCE’s activity upon solid PET ranges from poor to undetectable

Next, we investigated TTCE’s ability to degrade PET itself, using either PET granules, or PET granules turned into PET films through solubilisation of granules in HFIP, and evaporation of the HFIP. The film obtained from a defined weight of granule(s), i.e., ∼6 mg per granule, following evaporation of HFIP, was treated with 7.4 µM enzyme for 96 hours at 60 °C. Activity was assessed through the use of HPLC to separate species at the end of 24, 48, 72 and 96 hours, using separately set-up reactions. Figure 4D shows that with increasing duration of incubation, the amount of TPA obtained progressively increases, and also that the TPA produced after 96 hours is five times that produced after 24 hours. This shows that TTCE is able to act upon PET (although here it is after transformation of PET granules into films), with significant structural stability at 60 °C, over a 96-hour period. In stark contrast, when TTCE was directly added to PET granules that had not been turned into films through pre-processing, no TPA resulting from such incubation was detectable, as seen in Figure S6 †. Thus, it emerges that while TTCE acts upon PET chains organized into films of a few microns (in thickness), which present a large surface area, TTCE does not act substantively upon PET chains organized into a complex solid substrate. This essentially demonstrates that TTCE does not act upon PET, for all practical purposes, since it would not be expected to act upon post-consumer PET (as demonstrated in a later section). We further re-confirmed TTCE’s ability to degrade thin films (2 µM enzyme; 60 °C; 50 hours) by using TTCE upon commercially-sourced PET film discs (dia. 6 mm; 250 µm thickness) to produce detectable TPA and some BHET, as shown in Figure 4E, with the control PET film showing no significant hydrolysis under identical experimental conditions.

### TTCE and LCC work differently upon different PET-derived substrates

In this section, we present comparative data for the functioning of LCC, and TTCE, upon a number of different substrates: (i) PET films (whole PET chains), (ii) BHET (i.e., TPA flanked by two EG groups), (iii) para-nitrophenyl palmitate (a small aromatic phenyl group linked through an ester bond to a long aliphatic chain), (iv) fluorescein dilaurate (a large aromatic fluorescein group linked through two ester bonds to two long aliphatic laurate chains; one each flanking the fluorescein on either side), and (v) fluorescein di-benzoate (a large aromatic fluorescein group linked through two ester bonds to two aromatic benzoate groups, one each flanking the fluorescein on either side). The use of such diverse substrates in a comparative study facilitates comparisons of the activity-related characteristics of these two enzymes with high structural similarity, in respect of the different extents to which substrates of different grades of bulkiness (as well as different grades of hydrophobicity) tend to be acted upon by either enzyme, in a manner made even more interesting by the prior structural analyses reported in earlier sections, which showed that LCC and TTCE differ in respect of whether, or not, they possess (a) a deep catalytic cleft covered by structures that act like a lid upon the cleft (as is the case with TTCE), with limited hydrophobic surface area, or (b) a shallow catalytic cleft, surrounded by extensive hydrophobic surface area (as is the case with LCC).

#### TTCE is ∼2 times better at hydrolysing para-nitrophenyl palmitate than LCC

We first compared the activities of LCC and TTCE with the aliphatic long-chain ester, para-nitrophenyl palmitate, using identical reaction conditions for both enzymes (2 µM enzyme; 500 µM substrate; 70 °C; 5 hours). Figure 5A shows that TTCE is about twice as efficient, at hydrolysing palmitate ester, as LCC.

**Figure 5:**
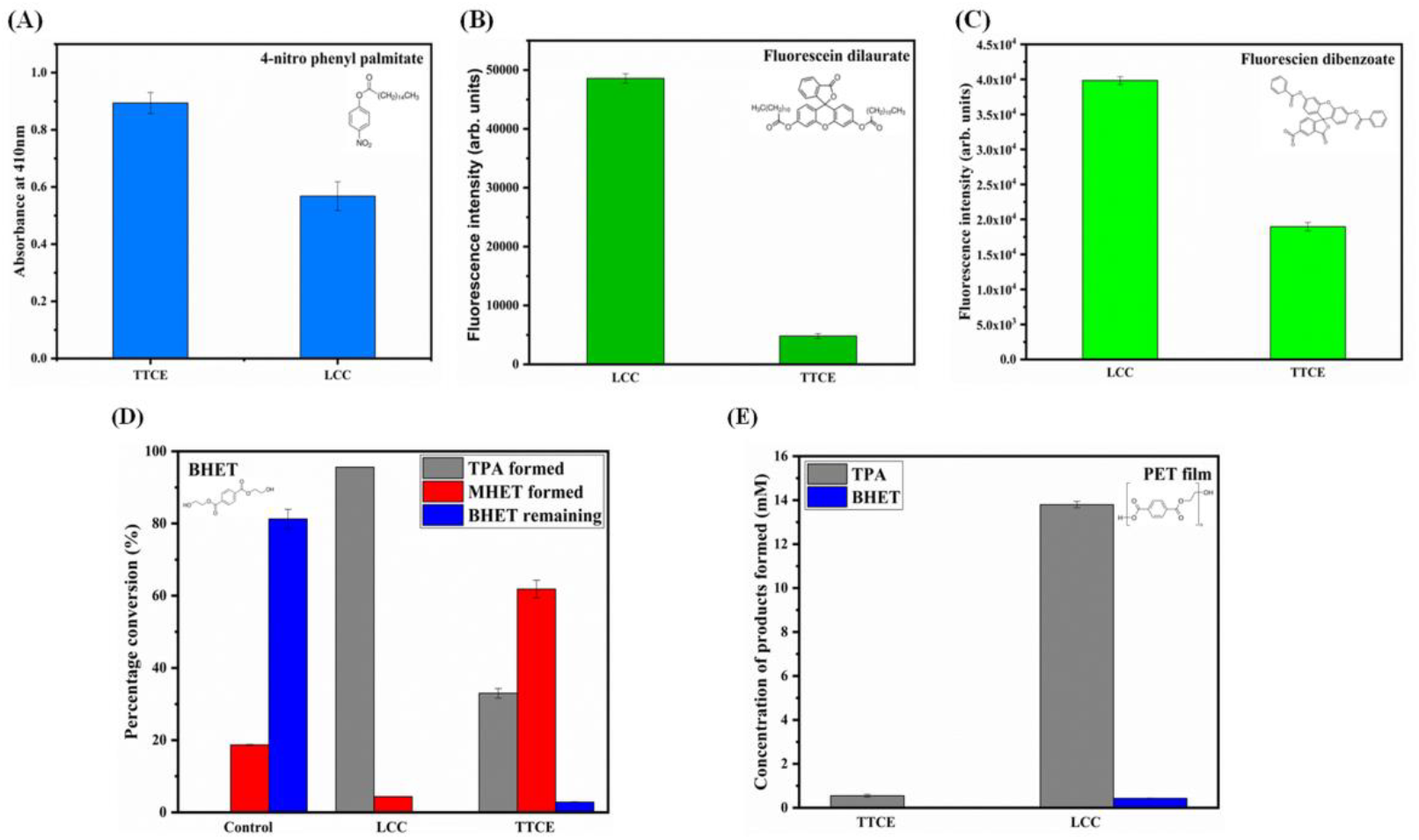
Comparison of the activities of TTCE and LCC upon (A) para-nitrophenyl palmitate. (B) Fluorescein dilaurate. (C) Fluorescein di benzoate. (D) BHET. (E) PET film.

#### LCC is ∼10 times better than TTCE at hydrolysing fluorescein dilaurate

In contrast, when similar experiments were performed with fluorescein dilaurate, as shown in Figure 5B, the activity of TTCE was found to be 10-fold lower than that of LCC.

#### LCC is only ∼2 times better than TTCE at hydrolysing fluorescein dibenzoate (comparable to BHET)

In contrast to both of the above experiments using different substrates of relatively small size, when a third such substrate, fluorescein dibenzoate was used in place of fluorescein dilaurate, as shown in Figure 5C, TTCE was found to perform only ∼2-fold poorer than LCC, and not ten times poorer (as was the case with fluorescein dilaurate).

#### LCC is ∼3 times better than TTCE at hydrolysing BHET

We next examined the activity of LCC and TTCE upon BHET, as shown in Figure 5D. BHET is a DI of PET, but in this experiment there was no PET present. The BHET was commercially-sourced. LCC was found to be efficient at turning BHET into TPA, leaving very little residue of MHET at the end of the period for which the reaction was performed. With TTCE, the residue of un-degraded MHET observed was larger, under the same conditions, suggesting that the conversion of BHET into TPA by TTCE is about ∼3-fold poorer than that carried out by LCC (34 % as compared to 100 % for LCC). In other words, in respect of enzymatic degradation of BHET, in the absence of any PET in the reaction environment, we found TTCE to be poorer than LCC, but only slightly poorer. Of course, in the presence of PET, as already shown, the bulk of the LCC happens to be titrated to the surface of PET, with consequences for the degradation of the PET-derived, LCC action-derived, BHET that escapes into solution.

#### LCC is ∼25 times better at hydrolysing PET than TTCE

Next, we assessed LCC and TTCE in respect of their abilities to hydrolyse PET, as shown in Figure 5E. For these experiments, PET was presented to both enzymes independently, in the form of commercially-sourced PET film (note: rather than the in-house generated PET film made using HFIP dissolution of PET granules, followed by evaporation of HFIP). Each of the two enzymes (at 2 µM concentration) was incubated with a single PET film of defined dimensions (6 mm diameter; 250 µm thickness) for a period of ∼50 hours, at 60 °C. The result obtained was that the amount of TPA generated from PET by LCC was found to be ∼25-fold higher than that generated by TTCE. Therefore, TTCE could be held to be incapable of degrading post-consumer (product-derived) PET, in comparison with LCC. Indeed, we also performed an experiment with post-consumer PET, and with PET granules that had not been turned into films (through HFIP dissolution), in order to test this contention. The data for this experiment is shown in Figure S6 †. The data shows that TTCE generates no detectable degradation products whatsoever, from either PET granules (Figure S6A †), or from post-consumer PET (Figure S6C †) taken from PET containers (bottles). On the other hand, LCC was clearly able to generate some detectable TPA (retention time ∼13.6 min in the HPLC traces shown in Figure S6) from both PET granules (Figure S6B †) and post-consumer PET (Figure S6D †).

### The synergistic action of LCC and TTCE improves generation of TPA from PET by ∼2.0 fold and generation of MHET by ∼2.5 fold

In this section, we examine whether the joint action of LCC and TTCE results in greater yields of TPA from PET than is obtained through the action of (i) LCC alone, (ii) TTCE alone, (iii) the sum of the mutually-independent actions of LCC and TTCE, or (iv) the sum of the actions of LCC and TTCE molecules that have been joined together in genetic fusion.

Figure 6 shows that the joint action of LCC and TTCE results in significantly greater yields of TPA and MHET from PET, with respect to what is obtained through LCC alone. Figure 6A demonstrates that the joint action of LCC and TTCE is significantly better than the sum of the individual actions of LCC and TTCE, when 100 nM TTCE, and 50 nM LCC, are used in a degradation experiment. The improvement in TPA generation is effectively by about ∼95 %, or a nearly ∼2-fold improvement, and the improvement in MHET generation is by about ∼147 %, or a nearly ∼2.5-fold improvement. Figure 6B shows that this improved joint action continues to be seen when 2 µM concentrations of LCC and TTCE are used. In this figure, it is also observed that the residual MHET (which was significant for all reactions when 100 nM TTCE and 50 nM LCC were used) can no longer be seen at the end of a reaction in which LCC and TTCE are acting synergistically with both enzymes at the higher concentration of 2 µM, although residual MHET is still seen with the reaction in which LCC is present at a concentration of 2 µM. These results establish both (i) that the joint action of LCC and TTCE significantly improves the yield of MHET, and TPA, from PET, and (ii) that this joint action also improves the conversion of the MHET that is generated by LCC into TPA. Additionally, data is also presented in Figure 6B for the action of a fusion protein consisting of an LCC domain and a TTCE domain, i.e., an N-terminal LCC domain joined to a C-terminal TTCE domain through a glycine- and serine-rich linker separating the two domains, (GGGSGGSGGGSGKLGGGSGGSGGGSG) with a 6xHis affinity tag at the C-terminus, following the TTCE domain. It is observed that the TPA yield from the fusion enzyme, marked ‘LCC-TTCE’, falls somewhere between the TPA yield obtained from LCC alone, and the TPA yield from the jointly (i.e., synergistically) acting combination of LCC and TTCE, marked ‘LCC+TTCE’.

**Figure 6:**
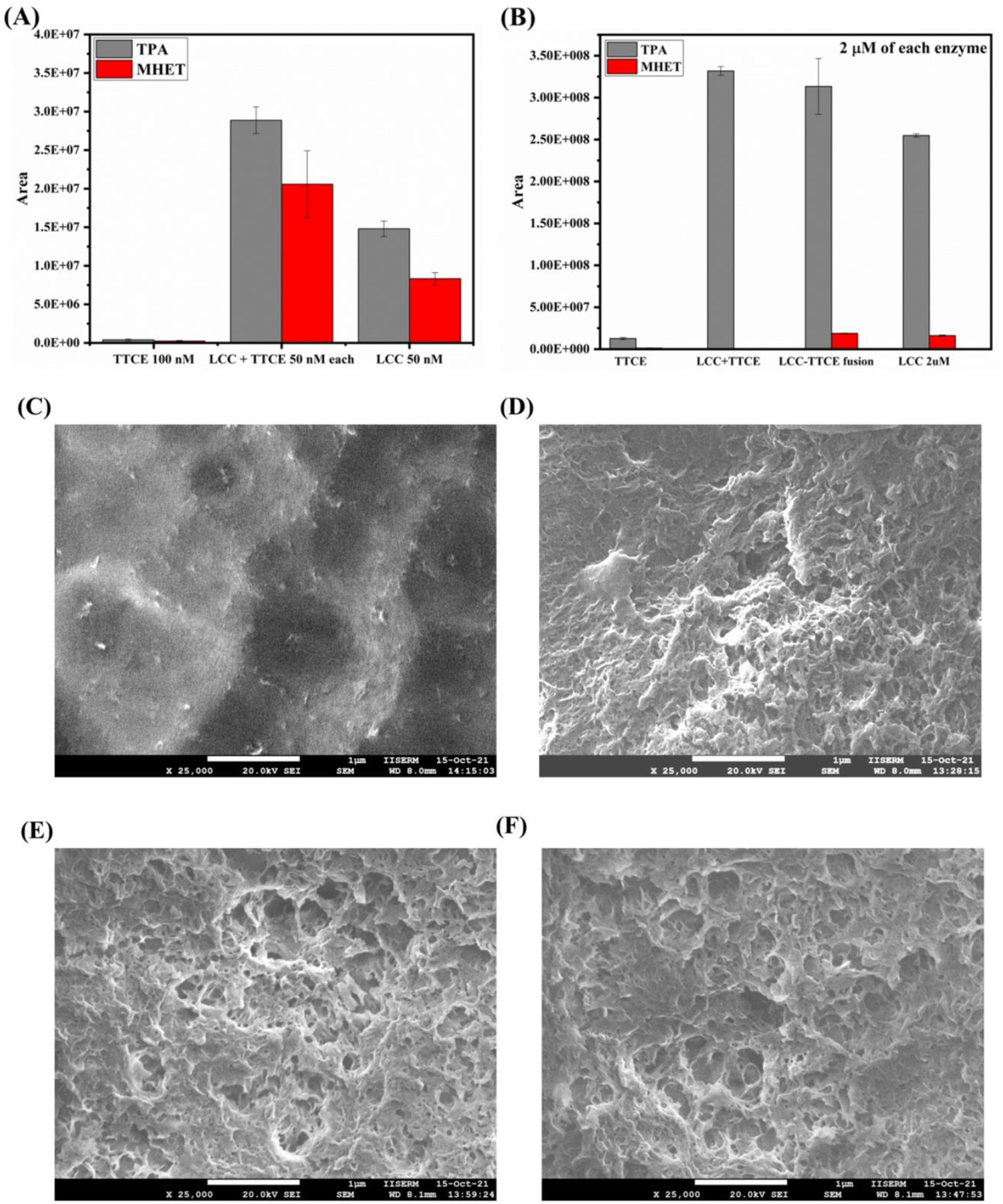
Synergy in the actions of LCC and TTCE. (A) Activities shown by 100 nM TTCE alone, 50 nM each of TTCE and LCC, and 50 nM LCC alone. (B) Activities shown by 2 µM TTCE alone, 2 µM each of TTCE and LCC, 2 µM of TTCE-LCC fusion construct, and 2 µM LCC alone. (C-F) Scanning electron microscopy (SEM) images of the surfaces of PET films subjected to exposure to LCC, the LCC-TTCE fusion construct, and the LCC+TTCE enzyme cocktail. Magnified images are presented. (C) Untreated PET surface (25000X). (D) Surface treated with LCC alone (25000X). (E) Surface treated with the LCC-TTCE enzyme fusion (25000X). (F) Surface treated with the TTCE+LCC enzyme cocktail (25000X).

It is also observed that the error bars for the experiment involving this ‘LCC-TTCE’ fusion are less tightly distributed around the mean value, than happens to be the case for the ‘LCC+TTCE’ experiment. We would like to rationalize this observation in the following manner. On the one hand, it is reasonable to suppose that the LCC-TTCE fusion would be largely titrated to the surface of PET like LCC alone, causing its action to be closer to that of LCC alone, rather than like that of LCC and TTCE acting jointly as ‘LCC+TTCE’. On the other hand, the SDS-PAGE data (see Figure S7 †) suggests that the glycine-serine-rich linker undergoes a significant amount of proteolytic cleavage during the prolonged incubation of 40 hours used for this binding assay, with this causing a certain amount of TTCE to also be present in solution, exactly as in the LCC+TTCE experiment, with this presumably leading to the LCC-TTCE fusion performing significantly better (rather than only somewhat better) than LCC alone, with the amount of TPA generated being closer to that obtained in the LCC+TTCE experiment. However, since this amount could vary from experiment to experiment, this explains the spread seen in the error bars for this data, compared to the other data which is characterized by much tighter error bars. Thus, in summary, although the LCC-TTCE fusion experiment could potentially be improved by making the linker less susceptible to proteolysis by contaminating proteases during the prolonged reaction, it is clear that a cocktail of LCC and TTCE (i.e., LCC+TTCE) performs better than a fusion of LCC and TTCE which, in turn, performs better than the sum of the individual activities (i.e., TPA yields) of LCC and TTCE working in each other’s absence, which in turn is better than the actions of either LCC alone, or TTCE alone, upon PET.

In Figure 6C-6D, we show qualitatively that ‘LCC+TTCE’ and ‘LCC-TTCE’ are better than LCC alone, with help of representative scanning electron micrograph (SEM) images shown at a magnification of 25000X. In the same figure, we also demonstrate that LCC invades PET surfaces locally, i.e., there are sites seen in the SEM images which show that some enzyme-treated surfaces have undergone excessive degradation in comparison to other neighbouring regions, as well as sites that appear to be largely unaffected by the enzyme. This suggests that the sites of original binding and localization of LCC molecules are also the sites at which these molecules largely remain, and continue to act. Therefore, it may be inferred that while LCC is three-times better than TTCE at degrading BHET when there is no PET present, LCC is unlikely to remain available in solution to degrade BHET when it is in the presence of PET, since it is likely to become titrated on to the surface of PET. In Figure S8 †, SEM images of higher magnification (75000X) are shown, in support of the same conclusions.

## Conclusions

In the work presented in this paper, we have introduced several advances in the area of PET degradation by enzymes: (1) We have argued that the improvement of PET invasion (and degradation) by a PET-degrading enzyme essentially works at cross-purposes with the need to improve the conversion of PET’s degradation intermediates into TPA, because the latter happen to be soluble (unlike PET itself), causing them to escape away from surface of PET (and any PET-bound enzymes), into solution, where they display a tendency to accumulate. (2) We have proposed that the solution to the problem of accumulation of degradation intermediates in solution could be the introduction of a second enzyme into the PET degradation reaction mixture; a soluble enzyme that can be driven to high concentrations, and made to work primarily to convert PET’s degradation intermediates into TPA, without engaging overly with the surface of PET. (3) We have identified such a second enzyme, or ‘assistant’ enzyme (TTCE), which is capable of working at, or above, the glass-transition temperatures of all forms of PET, for durations that are essentially unlimited, due to the enzyme’s high solubility, high thermal and chemical stability, and high kinetic structural stability. (4) We have shown that this ‘assistant’ enzyme engages poorly with PET, unlike a ‘master’ PET-invading enzyme (LCC), and also that it differs sufficiently from the master enzyme, in terms of the structural-biochemical features of its surface, and its active site, to prefer to degrade PET’s degradation intermediates over the degradation of PET itself. (5) We have shown that even though this assistant enzyme (TTCE) is unable to generate TPA from PET itself, it improves the generation of both MHET and TPA from PET when it is allowed to work synergistically with the master PET-degrading enzyme (LCC), essentially by engaging in a clear division of labour, in which one enzyme (LCC) degrades PET, and the other enzyme (TTCE) degrades the degradation intermediates of PET, to improve the yields of the terminal degradation intermediate (MHET) and the terminal degradation product (TPA). (6) We have also shown that, through the selection of suitable enzyme concentrations, the accumulation of residual MHET can be done away with altogether, through the effective use of the said division of labour between enzymes.

There is a reason that PET’s degradation intermediates are more soluble in water and aqueous media than PET itself. Both BHET, and MHET, have a higher content of polar hydroxyl groups per aromatic moiety (a ratio of 2:1) than PET chains, as can be seen in Figure S1 †. This causes BHET and MHET to have a higher likelihood of diffusing away from the solid PET substrate into the aqueous solution. Our contention in this paper has remained that both BHET, and MHET, which are released into aqueous solution, require that an enzyme that is capable of remaining in solution, and transforming them into TPA. Although saturation of the binding surface of PET by the master enzyme, LCC, could still theoretically allow a certain amount of LCC to remain in solution (to act upon BHET and MHET), the data in Figure 6 shows incontrovertibly that addition of TTCE clearly improves yields of MHET and TPA, under conditions involving non-saturation of PET’s surface by LCC. Since PET degradation continually exposes new PET surfaces, the saturation of PET’s surface by the master enzyme, LCC, is merely a theoretical possibility that is unlikely to ever be achieved in reality.

The structural bioinformatics-based analyses presented at the beginning of the paper predicted that the presence of the catalytic serine within a deep and narrow catalytic cleft in TTCE, as compared to LCC, as well as the presence of a ‘lid-like’ structure over TTCE’s active site must cause TTCE to accommodate much smaller substrates than LCC. We were able to establish this to be a fact, by reacting the two enzymes with a variety of substrates, and by showing that TTCE works more efficiently than LCC, or as efficiently as LCC, upon some of these substrates in proportion with the smallness of the size of these substrates. We were also able to show that TTCE works only somewhat poorer than LCC upon BHET, and upon fluorescein-dibenzoate [which may be considered to be a BHET analog of sorts^29^], as well as an analogue of short PET chains, or OET (consisting of 2 to 5 ester-linked MHET residues).

We have shown that the T_m_ of TTCE is higher than that of LCC, as assessed by DSC experiments involving both enzymes, as seen in Table S3 and Figure S9 † in the supporting information. The enthalpic component of TTCE’s thermodynamic stability (ΔH) is also higher than that of LCC. TTCE is also extremely kinetically-stable at high temperatures. Such high kinetic stability, associated with poor rates of unfolding, may be anticipated to allow TTCE to remain folded and functional in solution, for considerable lengths of time, at temperatures in the neighbourhood of its equilibrium T_m_ of 80.5 °C. Additionally, we have also presented data to demonstrate the extremely high chemical stability of TTCE to denaturation by Gdm.HCl (a C_m_ of 5.1 M) and urea (a C_m_ exceeding 8M, and expected to exceed the solubility of urea). This high chemical stability suggests that TTCE could be a useful enzyme in an industrial environment in which numerous additional chemicals and solvents could be present. Therefore, the stability properties of TTCE add to its value as an enzyme for the degradation of BHET, especially if TTCE were able to add value to PET degradation by working in a synergistic relationship with LCC.

In ending this paper, we request readers to also read Section 1 in the Supporting Information, which constitutes both an extended introduction, and an extended discussion, of all approaches to PET degradation (including those that are sustainable and environmentally-friendly), with a focus on all of the previous work that led to (and facilitated) the work presented here.

## Acknowledgements

We thank Dr. Arpana Jha for advice regarding the thermodynamic and kinetic analyses of TTCE. We thank the central facilities of IISER for access to MS-MS and SEM instruments, and the Technology Business Incubator at IISER Mohali for paid access to its HPLC facility. AM thanks the Department of Biotechnology, Government of India, for a doctoral research grant. BT thanks the Council of Scientific and Industrial Research, Government of India, for a doctoral research grant. PG thanks IISER for research funding, and also the Ministry of Human Resource Development, Government of India, for a Centre of Excellence grant to the CPSDE.

## Author contribution statement

AM and PG together conceived and designed ideas and experiments and wrote up the manuscript. AM performed all the experiments, and anchored the work. BT performed some of the experiments along with AM. AM and BT worked under PG’s supervision. The work is part of AM’s doctoral thesis.

## Conflict of Interest Statement

There are no conflicts to declare

## Data availability statement

Data is available from the authors on request. There is no separate file containing data, and all the data has been provided in the manuscript, or supporting information.

## Notes

### Competing Interest Statement

The authors have declared no competing interest.

