## Supporting Information for "Enhancing high-temperature degradation of polyethylene terephthalate through a synergistic division of enzyme labour between a solid-degrading thermostable cutinase and a reaction intermediate-degrading thermostable carboxylesterase"

Running Title: *A dissolved carboxylesterase aiding a plastic-bound cutinase*

* Corresponding Author

Webpage: www.guptasarmalab.in

#### Contents

**Section 1. Green chemistry-related aspects of this work (Pages 3-8)**

**Section 2. Materials and methods (Pages 9-13)**

**Section 3. Tables (Page 14)**

**Table S1:** Docking and binding scores.

**Table S2:** Basic properties of TTCE, LCC, TTCE-LCC fusion construct.

**Table S3:** Thermal, kinetic and thermodynamic stabilities of TTCE and LCC.

**Section 4. Figures (Pages 15-23)**

**Figure S1**: Enzymatic breakdown of polyethylene terephthalate (PET)

**Figure S2**: RMSD and RMSF from molecular dynamics simulation of TTCE-2HE-$(MHET)_4$.

**Figure S3:** Biophysical characterization of TTCE.

**Figure S4:** Thermal and chemical stability of TTCE.

**Figure S5:** Chemical kinetic stability studies of TTCE.

**Figure S6**: Degradation of intact PET granules and post-consumer PET by TTCE and LCC.

**Figure S7**: The PET-film bound fraction of the LCC-TTCE fusion construct.

**Figure S8**: Scanning electron microscopy (SEM) images of the surfaces of PET films subjected to enzymatic treatments.

**Figure S9:** Thermal stability of LCC.

**Section 5. References (Pages 24-26)**

### Section 1. Green chemistry related aspects of this work

#### Processing of post-consumer PET for degradation:

#### A summary of approaches used

1) **Mechanical treatment.** PET is turned into micron-sized beads through a process of crushing, rolling and milling.[1]
2) **Heat treatment.** PET is decomposed either through pyrolysis (~400-600 ˚C), or through microwave treatment (160-250 ˚C) into terephthalic acid (TPA) and ethylene glycol (EG).[2]
3) **Thermo-chemical treatment.** A combination of chemicals and high temperatures is used to decompose PET. For example, (i) methanol treatment at 250-290 ˚C turns PET into dimethyl terephthalate (DMT) and ethylene glycol (EG), or (ii) treatment at 190 ˚C in ionic liquids such as supercritical ethanol with liquid 1-butyl, 3-methyl imidazolium tetrafluoroborate, or in the presence of the catalyst, zinc acetate, turns PET into bis-hydroxyethyl terephthalate (BHET).[2]

   The above methods have recently been reviewed.[2]
   [Damayanti and Ho-Shing Wu, *Polymers*, (2021) **13**, 1475-1512].

4) **Microbial/enzymatic treatment**. Microbe-secreted esterases act upon PET and the degradation intermediate, mono-hydroxyethyl terephthalate (MHET) is taken up by some fungi, actinomycetes, or preteobacteria (e.g., the organism, *Ideonella sakaiensis*). Inside the cell, other esterases turn MHET into protocatechuate (PCA) which enters the TCA cycle. This enables some microbes to degrade PET and use some degradation products as a source of carbon.[3]
5) **Pure enzymatic treatment.** Various esterase enzymes are currently being studied: (i) a cutinase, TfCut2, from *Thermobifida fusca*;[3] (ii) a cutinase, LCC,[3] derived from a metagenomic library created from leaf branch compost; (iii) a fusion enzyme consisting of a PET hydrolyzing esterase and a MHET-hydrolyzing esterase.[4] Some fungus-derived enzymes are also being explored.[5] Such enzymes bind to, and invade, solid crystalline PET at moderately high temperatures, to degrade it into a mixture of oligoethylene terephthalate (OET), BHET, MHET, TPA and EG.[5]

   The above methods have recently been reviewed.[3]
   [D. Danso, J. Chow, W.R. Streit, Plastics: environmental and biotechnological perspectives on microbial degradation. *Applied and Environmental Microbiology*, 2019, **85**:e01095-19].

#### PET enzymatic degradation/recycling:

#### A summary of (green chemistry-related) advantages

1) **Reduction in energy expenditure.** Use of lower temperatures. Cutinases/esterases from mesophile organisms (growth optima in the range of 20-60 ˚C) work optimally at temperatures in the range of 30-45 ˚C, and cutinase/esterases from thermophile organisms (growth optima in the range of 60-80 ˚C) work optimally at temperatures in the range of 50-70 ˚C.[6] Thus, enzymes obviate the need to use high temperatures (> 400 ˚C for thermal decomposition of PET, and > 200 ˚C for thermo-chemical decomposition). It is sufficient to achieve temperatures approaching ~ 60-80 ˚C, to cause PET to undergo a glass-transition and expose the polymer backbone to enzymes. No need for stirring. Enzymes also obviate the need to use energy to stir reaction mixtures.[7]
2) **Reduction in cost of materials used.** Exploitation of microbes as low-cost enzyme-producing factories. Thermostable PET-degrading enzymes are produced as recombinant enzymes, through biosynthesis, i.e., through heterologous expression in mesophile organisms that grow upon low-cost nutritional substrates, at low temperatures,[8] with each cell producing the desired enzyme at a

small fraction of the cost of producing the enzymes through chemical synthesis from amino acids.[9] With an enzyme such as LCC, using 3 milligrams of which, i.e., 0.3 % (w/w) 1 gram of PET can be almost fully degraded,[10] it is easy to produce 80-100 milligrams per litre of fermented *E. coli* culture, since 6-8 milligrams can be produced per litre of shake-flask cultures that allow one-fifteenth to one-twentieth of the growth achievable through high-density fermentation). With other enzymes, e.g., the TTCE enzyme used for the synergy demonstrated in this paper, the yields can be as much as four times this amount, in milligrams. *Exploitation of enzyme thermostability for low-cost purification.* The purification of recombinant thermostable enzymes produced by mesophile genetically-engineered microbial cell factories is of low cost due to the fact that cells can be heated to 60-70 ˚C to directly obtain enzyme of high purity, through thermal rupture of cells and denaturation and aggregation/precipitation of all cellular protein/enzyme constituents other than the desired (thermostable) enzyme, thus obviating the need for purification of enzymes by expensive chromatographic methods.[11]

3) **Elimination of toxic/expensive chemicals and solvents.** Unlike chemical methods, enzymatic methods use aqueous environments, and require neither the presence of any expensive/toxic chemicals, nor the use of any expensive/toxic solvents.
4) **Enhanced scope for recycling.** Degradation of post-consumer PET by thermal and thermo-chemical means mostly results in the production of heterogenous mixtures of TPA and other degradation intermediates (including side-products that are colored), although some expensive methods that are not commercially-viable do result in very high yields of TPA approaching 80-90 %.[12] The presence of side-products along with PET degradation intermediates in the TPA obtained through processing of post-consumer PET thus adversely affects the cost of production, and also the quality, of the PET that can be produced through recycling of such 'reclaimed' TPA.[12] In contrast, with enzymatic degradation, there are no side-products. If the yield of TPA from enzymatic degradation of PET can be improved to nearly one hundred percent, the PET that would be generated from such TPA is expected to be of comparable quality to virgin PET.[10]
5) **Reduced consumption of fossil fuel (petroleum).** PET is currently made from petroleum.[13] Enzymatic degradation of PET into TPA which can be recycled into food-grade (virgin-like) PET is likely to reduce dependence on fossil fuel (petroleum) for the production of PET of acceptable quality through recycling, thus creating a viable circular economy involving PET and TPA that would reduce consumption of petroleum. Notably, it has been shown that the cost of production of new PET plastic from TPA generated through enzymatic degradation of post-consumer PET is about 4 % of the cost of production of fresh PET from petroleum.[10]
6) **Enhanced scope for valorization.** The TPA generated from PET through enzymatic degradation is of high-enough quality for it to also be turned into high-value aromatic products, or aromatic-derived products, such as protocatechuic acid (PCA), gallic acid (GA), pyrogallol, catechol, muconic acid (MA), and vanillic acid (VA).[14,15] These are used in the manufacture of pharmaceuticals, cosmetics, sanitizers, animal feeds, bioplastic monomers, and so on.[14]

#### PET enzymatic degradation/recycling:

#### A summary of issues and previous attempts at resolution

***Issues***

1) **The generation of non-TPA intermediates, and the cost of separating TPA from these intermediates.** The main thing that appears to be currently holding back the recycling of PET in order to create a circular economy (with reduced dependence on fossil fuels, and increased prospects for valorization of TPA) is that enzymatic degradation of PET can occur upon a variety of different ester bonds in PET, and this causes the early stages of enzymatic degradation to inevitably produce not just pure TPA (which is the end product that is expected to result from the

breakage of all ester bonds in the PET backbone), but rather a mixture of oligoethylene terephthalate (OET), BHET, MHET, TPA and EG instead. Without separating TPA away from these other (intermediate) products of PET degradation, recycling of TPA into PET remains unviable, because separation is a costly proposition.[16,17] What is needed is an enzymatic degradation system that leaves no intermediates. For this, it is necessary to pay attention to multiple possible ways in which such intermediates fail to be hydrolyzed to a significant degree.

2) **The inhibitory effect of non-TPA intermediates.** Certain degradation intermediates such as MHET appear to inhibit the PET-hydrolyzing activity of enzymes. In particular, TfCut2 displays significant MHET-based inhibition.[18] Notably, LCC also shows MHET-based inhibition, although this is to a far lower degree than seen with TfCut2. Due to this, it is necessary to pay attention to multiple possible ways in which to either neutralize, overcome, or go around the problem of MHET-based inhibition.[19]

***Attempts at resolving issues, and their outcomes***

1) **Improvement of the PET-hydrolyzing activities of cutinase/esterase enzymes.** *Improvement of catalytic rates.* One approach that has been taken is to improve the enzymes that degrade PET, perhaps hoping that this would lead to more comprehensive degradation of both PET and its degradation intermediates. There is no doubt that this approach, involving both (a) the search for better enzymes, and (b) the improving of the activities and stabilities of such enzymes, through rational or combinatorial protein engineering, or other approaches, such as enzyme immobilization, has resulted in improvements in PET degradation. The search for better enzymes has led to the identification of TfCut2 from *Thermobifida fusca*, and LCC from a leaf branch compost metagenomic library.[6] These are currently the leading naturally-occurring enzymes that have been identified, with LCC being clearly being more efficacious than TfCut2.[10] Attempts have been made to improve both enzymes through protein engineering, and the current leading enzymes are variants of TfCut2 and LCC.[10,19-22] Thus far, these improvements have managed to create a variant of LCC that is ~27 % more active (i.e., 1.27-fold more active) than LCC.[10] *Improvement of enzyme stability - leading to longer timescales of activity, and to improved catalytic output.* Attempts have been made to improve the thermal stabilities of enzymes, with a view to increasing their longevity in reactions (and, therefore, their overall activity, as a function of the durations of reactions).[10] *Improvement of PET binding leading to greater residence-times of enzymes upon solid PET, and to improved catalytic output.*[23,24] Attempts have also been made to improve the PET-binding affinities of enzymes, with a view to increasing the residence time of enzymes upon PET, in order to improve the overall activity.

   **Outcome:** Despite improvements in yields of TPA resulting from improvements in enzyme kinetic rates, enzyme thermal stabilities, and enzyme binding to PET, the generation of mixtures of degradation intermediates along with TPA remains a persistent issue. It has thus far not proved to be possible to generate pure TPA through enzymatic reactions. Basically, although it has been proved to be possible to improve the yields of TPA, this improvement has resulted in the concomitant improvement of the yields of the other intermediate degradation products as well.[18,25] Thus, enzyme improvements have increased the production of TPA, but not improved the quality/purity of the generated TPA.

2) **Use of enzyme synergy, to reduce MHET inhibition.** It has been perceived that it could be useful to deploy a dual-enzyme system, i.e., to have an additional enzyme present along with cutinase engaged in PET hydrolysis (e.g., TfCut2, or LCC), to use this additional enzyme to reduce the amount of MHET, since MHET inhibits TfCut2 to a significant degree, and also inhibits LCC

(although to a lesser degree).[18]Thus, an immobilized carboxylesterase has been used along with cutinase/esterase (either TfCut2, or LCC), using a fixed 10 µg/ml concentrations of the cutinase/esterase, and varying concentrations (0-30 µg/ml) of an immobilized carboxylesterase.[18]

**Outcome:** Use of the specific dual-enzyme system led to (A) a significant improvement in the ratio of MHET with respect to TPA (i.e., the ratio was reduced), as desired. However, (B) significant levels of MHET were still present. Further, (C) perhaps owing to the reduced MHET-based inhibition, TPA and MHET levels both increased.

Therefore, enzymatic degradation has thus far not succeeded in making the recycling of TPA generated from post-consumer PET back into PET viable, mainly because the TPA that is generated exists in a mixture with other degradation intermediates such as OET, BHET and MHET. So, although enzymatic degradation remains a desirable goal for every one of the reasons listed in the previous section (from the viewpoint of Green Chemistry), and also because it produces no side-products that cannot be resolved into TPA upon further esterase treatment, the goal has not yet been achieved.

#### PET enzymatic degradation/recycling:

#### A summary of the philosophy and approach of the new proposal that is presented in this work.

1) **Making improvements in PET-hydrolyzing cutinases/esterases alone worsens the problem.** Cutinases that hydrolyze PET are required to bind to solid PET. In proportion with their ability to bind to solid PET, therefore, such enzymes happen to become titrated onto the surface of PET. This titration naturally depletes the enzyme population that is available in aqueous solution.[23]

2) **Depletion of enzyme from solution (due to enzyme binding to PET) is the main problem.** We argue that it is this depletion that underlies the fact that mixtures of TPA and other degradation intermediates are always obtained, from every PET hydrolysis reaction, regardless of how efficient an enzyme is (at hydrolyzing PET and its degradation intermediates which have the same polymer backbone chemistry as PET).[18,25]

3) **Cutinases/esterases are actually more efficient at degrading the degradation intermediates than they are at degrading PET, but degradation intermediates still accumulate.** It is a fact that is known to all who have worked with these enzymes that cutinases such as TfCut2, and LCC, efficiently generate TPA from both BHET and MHET, but also that the very same enzymes generate large amounts of OET/BHET/MHET that remain unresolved into TPA, as residues in solution, when they act upon PET, without degrading these into TPA (despite being able to do so more efficiently than they are able to degrade PET).[18,23,25] It is a known fact that PET hydrolysis reactions can have TPA and MHET present in a ratio of 60:40, at the end of a long incubation that lasts a few days.

4) **Cutinases/esterases are actually capable, but unavailable, to degrade the degradation intermediates into TPA.** Accumulation of OET/BHET/MHET should ideally not have occurred if PET-hydrolyzing cutinases/esterases happen to be both able to degrade MHET more efficiently than PET, and also available to do so, during PET hydrolysis. We point out that these enzymes are indeed able to degrade OET, BHET and MHET (which are all less hydrophobic than PET, and sparingly soluble in water, and which can diffuse into and out of enzyme active sites) more

efficiently than they are able to degrade PET[26,27] (which is hydrophobic and consists of densely packed chains of polymers that are insoluble). Even so, because these enzymes bind to PET, owing to their surface hydrophobicity, and then remain preoccupied with PET, they do not remain available in solution to degrade the degradation intermediates and the TPA molecules that escape into solution, when an enzyme performs a hydrolysis reaction on the surface of solid PET.[23,24] We argue, therefore, that the titration of the cutinase/esterase enzyme population onto the surface of PET is the primary reason for the accumulation of residual and unresolved OET/BHET/MHET, since there is no enzyme available in solution to resolve OET, BHET and MHET into TPA. We propose that this situation persists for as long as there is any solid (undegraded) PET still left (and available) to titrate away all hydrolyzing enzyme molecules away from solution. We argue that this situation is only likely to get worse with the further improvement of the abilities of different enzymes to bind to PET and hydrolyze it, if attention is not paid to placing enzymes in solution for the specific purpose of hydrolyzing OET, BHET and MHET into TPA.

5) **The solution probably lies in the making available of a second enzyme; one that is specifically deployed in solution.** Thus, we argue that the only way of resolving the OET, BHET and MHET that accumulates in the solution around solid PET is to make a different enzyme available for degrading these molecules, in concert with the action of the cutinase/esterase upon solid PET, i.e., a second enzyme that does not either bind to, or hydrolyze, PET but which is able to degrade OET, BHET and MHET.

6) **A two-fold division of labor.** The two enzymes of a dual-enzyme system would ideally engage in a two-fold division of labor; one in terms of the chemicals that the two enzymes work upon, i.e., PET, on the one hand, and PET's degradation intermediates, on the other hand; and the other in terms of the locations at which the two enzymes perform their work, i.e., one on the surface of solid PET, and the other in the solution around solid PET. With such a division of labor, neither enzyme would interfere either physically, or chemically, with the work being performed by the other enzyme.

7) **Release of MHET-based inhibition as a bonus, rather than as a goal.** It may be noted that if an enzyme degrades MHET in the solution around PET, the levels of MHET in the vicinity of PET's surface would reduce, thus relaxing the MHET-based inhibition of cutinase/esterases working at PET's surface. Thus, the need to resolve OET, BHET and MHET into TPA, in the solution around PET, in order to improve the quality of TPA, and the need to release the inhibition of PET-hydrolyzing enzymes by MHET, are both served by the same approach, namely to deploy a OET/BHET/MHET hydrolyzing enzyme in the solution around PET, so that the degradation intermediates of PET are further resolved into TPA concomitantly with their generation.

8) **Carboxylesterases are probably best suited to playing the role of 'second' enzyme.** Carboxylesterases are already known to degrade MHET.[18] We think that carboxylesterases are ideally suited to degrading small aliphatic as well as aromatic and other esters, because the folds of the polypeptide backbones are somewhat homologous to those of cutinases/esterases, but their active sites tend to be much deeper and less hydrophobic that those of PET-hydrolyzing enzymes, causing carboxylesterases to be incapable of binding to PET (due to PET being unable to access the catalytic residues which are buried away deep with the active site, and also due to PET being too hydrophobic to bind to an active site that is more hydrophilic than the active sites of cutinases/esterases.

#### PET enzymatic degradation/recycling:

#### A summary of what we have achieved using the approach outlined above.

As explained above, enzymatic degradation of PET comes with many benefits from the viewpoint of Green Chemistry.

Much prior work in this area has been done. Many excellent enzymes have been discovered, and it may be anticipated that more such enzymes will be discovered, in due course.

The main issue that remains, however, is the incomplete degradation of PET, with the generation of degradation intermediates that remain in reaction mixtures as residues.

Our arguments and proposals are in favor of the possible complete degradation of PET through the use of an additional enzyme.

We demonstrate that the complete degradation of PET can be achieved by using an additional enzyme that fulfills multiple criteria.

(1) Demonstrated extreme thermostability (demonstrated in this paper),
(2) Demonstrated extreme solubility (demonstrated in this paper),
(3) Demonstrated extreme producibility (demonstrated in this paper),
(4) Demonstrated extreme substrate versatility (demonstrated in this paper),
(5) Demonstrated inability to bind to PET (demonstrated in this paper),
(6) Demonstrated inability to perform any significant hydrolysis of PET (demonstrated in this paper),
(7) Demonstrated ability to work with the cutinase, LCC (demonstrated in this paper), and
(8) Demonstrated ability to outperform LCC, when working in concert with LCC, yielding the highest yields of TPA ever shown, and also the lowest accumulation of degradation intermediates ever shown.

Our approach, reasoning and results point to a possible way forward, and help to consolidate views that favor the use of dual-enzyme systems that involve a division of both physical (locational) and chemical (catalytic) labor.

### Section 2. Materials and methods

***Molecular docking and MD Simulation.***

Molecular docking and MD simulation studies were performed to determine residues likely to be involved in OET binding and hydrolysis in TTCE. The structure file for TTCE (RCSB PDB ID: 1UFO) was subjected to the Protein Preparation Wizard[28] to obtain a reliable all-atom structure. As the ligand used for docking, the initial structure file for an oligo-ethylene terephthalate (OET) chain of four MHET residues [2HE-(MHET)$_4$] was prepared using 2D Sketcher. Energy minimization was performed using the Schrödinger LigPrep module. Docking studies were performed using the Schrödinger Glide module,[29] with XP-Dock scoring function. The receptor grid was generated to create a 3D space in which the ligand was docked. Using XP-visualizer, docking scores and penalties were determined as detailed in Table S1. The Schrödinger Prime MMGBSA module[30] was employed to more accurately calculate the binding affinity of 2HE-(MHET)$_4$ to TTCE. The Schrödinger Desmond package was employed to perform MD simulations, using docked structures as the initial structures for the simulation. Desmond's System Builder tool was employed to prepare the system, and the OPLS3e force-field was applied, with simulation carried out in an orthorhombic box filled with TIP3P water as solvent. The system was neutralized through addition of counter ions and salt ions (concentration maintained at 150 mM NaCl). The Desmond MD simulation tool was used to carry out 75 ns of simulation, using a recording interval of 10 ps, within NPT ensemble at 300 K, using the Nose–Hoover chain thermostat,[31] and 1.0013 bars, using an integration time-step of 2 fs, and the Martyna–Tobias–Klein barostat.[32] A default cut-off radius of 9.0 Å was specified for Coulombic interactions. MD simulation trajectories were analyzed using Desmond's simulation interaction diagram (SID) tool. The free energy of binding ($\Delta G_{bind}$) was calculated using the OPLS3e force-field and VSGB 2.1 solvation model, through application of the following equation: $\Delta G_{bind} = E_{Complex} - E_{Ligand} - E_{Receptor}$, in which E is the energy associated with the van der waals, hydrophobic and electrostatic interactions in the ligand-protein complex ($E_{Complex}$), during ligand desolvation ($E_{Ligand}$) and during receptor desolvation ($E_{Receptor}$), respectively.

***Gene cloning, expression, purification and identity-confirmation of TTCE, LCC and LCC-TTCE fusion.***

The gene encoding TTCE was cloned from the genomic DNA of *Thermus thermophilus* into a pET23a plasmid, between Nde I and Hind III restriction sites, using the following primers to introduce restriction sites for cloning: NdeI-TTCE Forward : 5' TATATACATATGAGGGTTCGGACCGAGCGGCTC 3'. TTCE-HindIII : 5' ATATATAAGCTTCCGTGCCTCAAGCCAGTG 3'. The pET23a plasmid containing the TTCE gene was transformed into the BL21pLysS* expression host, induced to express TTCE with a C-terminal 6xHis affinity tag, using 1 mM IPTG (Isopropyl β-d-1-thiogalactopyranoside) at a culture optical density of 0.6, and induction for 5.5 hours at 37 °C following which cells were harvested through centrifugation, and the pellet obtained dissolved in bacterial cell lysis buffer (50 mM sodium dihydrogen phosphate, 150 mM NaCl and 10 mM imidazole) and subjected to sonication in the presence of lysozyme, before further centrifugation at 12,000 rpm to settle cell debris. The lysate was loaded onto a pre-equilibrated Ni-NTA (Nickel-nitrilotriacetic acid) column, for IMAC affinity chromatographic purification, using a wash with 35 mM imidazole, protein elution with 250 mM imidazole, and collection of protein in seven 1 ml fractions, followed by SDS-PAGE analysis. The gene encoding LCC (Genbank: AEV21261) was codon optimized for expression in *Escherichia coli*, gotten synthesized by a commercial service-provider (Biotech Desk Pvt. Ltd., Hyderabad, India), cloned without the gene segment encoding the signal peptide (between the Bam HI and Hind III restriction sites of the pQE30 vector) to produce a construct possessing an N-terminal 6xHis polyhistidine tag, and then expressed and purified using IMAC affinity chromatography from XL1Blue cells, using standard protocols. The gene encoding the LCC-TTCE fusion, incorporating a 22 residues-long polypeptide segment $(GGGSGGSGGSG)_2$ as linker between the LCC and TTCE segments, was created in two steps (through PCR reactions) and cloned between the Nde I and Not I restriction sites of pET23a, to produce (and purify, using IMAC affinity chromatography) a C-terminally 6xHis-tagged LCC-TTCE fusion, using the BL21pLysS* strain of *E. coli*. The gel band corresponding to a protein of the expected size was excised, subjected to trypsin treatment using the ProteoProfile™ Trypsin In-Gel Digest Kit (Sigma Aldrich, Product Code PP 0100), mixed with the MALDI matrix, α-Cyano-4-hydroxycinnamic acid (Sigma Aldrich, Product Code C8982) in a ratio of 1:1, and subjected to MALDI-Q-TOF analyses on a Synapt G2S-HDMS mass spectrometer (Waters), in single-stage (MS) mode, followed by matching of masses of observed peptides with the masses of peptides generated through *in silico* trypsin digestion of TTCE (using ProtParam and the ExPASy server). The basic properties of the three recombinant proteins produced and studied are detailed in Table S2.

***Spectroscopic and chromatographic characterization of the folded state of TTCE.***

*(1) Quaternary structure.* Size exclusion chromatography (SEC), glutaraldehyde crosslinking, and dynamic light scattering (DLS) were used to examine the oligomeric status of TTCE. *Firstly*, following IMAC chromatography, eluted samples were concentrated to a volume of 500 µl and loaded onto a Superdex-75 Increase 10/300 GL gel filtration column (GE-healthcare), pre-equilibrated with 25 mM sodium dihydrogen phosphate buffer of pH 8.0, on an AKTA workstation, to compare the elution volume of TTCE with a calibration standard, using SEC. *Secondly,* to check for the presence of higher order oligomers of TTCE, protein purified through SEC was incubated with 0.05%, 0.1% and 0.2% (v/v) of glutaraldehyde, for 10 minutes at room temperature, with stopping of the reaction through boiling of samples in dye-containing, 5x SDS-PAGE loading buffer, at 99 °C, for 5 minutes, followed by electrophoretic analyses of samples on a 13% acrylamide SDS-PAGE, with analyses of gels based on the known preservation of glutaraldehyde crosslinks during SDS-treatment and boiling, and observations of gel band mobilities. *Thirdly*, dynamic light scattering (DLS) based characterization was done in phosphate-buffered saline at pH 8.0 on a Wyatt QELS+ Heleos 8 instrument.

*(2) Tertiary structure.* The tertiary structure of TTCE was examined through assessment of its folded state based on fluorescence spectroscopic examination of the wavelength of maximal fluorescence emission of its intrinsic fluorescence, derived from its single tryptophan residue. The occurrence of a wavelength of maximal emission that was significantly lower than ~353 nm was used as evidence of the shielding of the tryptophan residue from the aqueous solvent through the formation of a tertiary structure.

(3) *Secondary structure.* The secondary structure of TTCE was examined on a Chirascan™ circular dichroism spectrometer (Applied Photophysics Ltd.), using a quartz cuvette of 1 mm path length, TTCE at a concentration of 0.2 mg/ml in 25 mM phosphate buffer of pH 8.0, and mean residue ellipticity (MRE) calculated at each wavelength, to create the CD spectrum, using the formula: MRE = (θ×mean residue weight×100)/(1000×concentration in mg/ml×pathlength in cm), where $\theta$ was the raw ellipticity measured in millidegrees.

***Spectroscopic and chromatographic characterization of the equilibrium thermal stability of TTCE's folded state.***

*Firstly*, thermal stability was assessed using circular dichroism (CD) through the thermal denaturation of TTCE (by heating between 20 °C to 90 °C, at a rate of 1°C/min) with concomitant collection of CD spectra on a Chirascan™ (Applied photophysics) spectrometer fitted with a Peltier block, using a protein concentration of 0.2 mg/ml in 25 mM phosphate buffer, pH 8.0, in a 2 mm path length (quartz) cuvette.

*Secondly,* thermal stability was assessed using differential scanning calorimetry (DSC) through the thermal denaturation of TTCE (heating between 20 °C and 90 °C, at a rate of 90 °C/h, and cooling between 90 °C and 20 °C, at a rate of 60 °C/h) with concomitant measurement of enthalpic changes on a differential scanning calorimeter (MicroCal VP-DSC), using a protein concentration of 0.5 mg/ml, following 5 cycles of heating and cooling of the control (phosphate) buffer, to generate the baseline, and data concerning heat required to unfold the protein being extracted from the system and measured, over the temperature range in which a difference in the rate of change of temperature between sample and control (reference) cells could be observed. A specific heat capacity *versus* temperature graph was obtained, and the area under the baseline-subtracted, concentration-normalized graphs was then estimated to measure the change in enthalpy associated with protein unfolding, with the peak of the transition curve assessed to be the measured melting temperature.

*Thirdly,* thermal stability was assessed using fluorescence spectroscopy through the thermal denaturation of TTCE (heating between 20 °C and 90 °C) with concomitant measurement of changes in the tertiary structure of TTCE through monitoring of red shifts in TTCE's fluorescence emission maximum wavelength, on a steady-state fluorimeter (FluoroMax, HORIBA), during thermal unfolding, with sample excitation performed at 295 nm, and emission recorded between 300 and 400 nm.

***Spectroscopic characterization of the chemical stability of TTCE's folded state.***

(1) *Equilibrium measurements.* Chemical stability was assessed using CD and fluorescence spectroscopy, for samples of TTCE incubated overnight with the chemical denaturants, urea, and guanidium hydrochloride (Gdm.HCl), using the monitoring of changes in spectral characteristics (i.e., CD spectral MRE values at 222 nm, to monitor changes in TTCE's secondary structure; and fluorescence emission spectral maximum wavelengths, to monitor changes in TTCE's tertiary structure) as a function of the unfolding of 0.2 mg/ml TTCE, in the

range of 0-8 M urea, or 0-6 M Gdm.HCl. Values of $C_m$ (i.e., the denaturant concentration at which half the molecules in any population have undergone unfolding) were determined for unfolding by the two denaturants.

*(2) Kinetic measurements.* To study the kinetics of unfolding, TTCE was incubated with Gdm.HCl (different concentrations in the range of 5-7 M) for two hours, with monitoring of changes in secondary structure as a function of time, and with calculation of the fraction unfolded with respect to time of incubation (2 hours) thereafter being plotted and fitted, using the 'expDecay1' function of the software, Origin Pro 2018. The respective rates of unfolding ($K_u$) determined from the plots were utilized to obtain the half-chevron plot (natural logarithm of $K_u$ *versus* activity of denaturant [D] determined from the concentration of denaturant [Gdm.HCl]). The slope of the half- chevron plot was used to calculate the rate of unfolding in absence of denaturant ($K_{u;w}$). This rate indicates the kinetic stability (i.e., resistance to unfolding) at a particular temperature. The relevant equations used were (1) $K_u = K_{u;w} \cdot e^{m_u \cdot D}$, in which $K_{u;w}$ is the rate of unfolding in absence of denaturant; and (2) D=7.5[Gdm.HCl]/(7.5+[Gdm.HCl]), in which D is the activity of denaturant and $m_u$ is the slope of half chevron plot.[33]

***Activity of LCC and TTCE upon different esters.***

*(1) Absorption measurements using turbidogenic or chromogenic substrates.* To assess TTCE's ability to act upon short and long chain aliphatic esters, the enzyme was reacted with 1-Naphthyl butyrate (2.5 mM), and 4-Nitrophenylpalmitate (250 µM), at temperatures ranging from 50 °C to 100 °C, releasing napthol and para-nitrophenol, which were quantified using absorbance measurements at 235 nm, and 410 nm, respectively. The catalytic potential of LCC in comparison with TTCE (2 µM each) was compared using the following two substrates: (i) 4-Nitrophenylpalmitate (250 µM) at 70 °C for 5 hours, and (ii) Fluorescein dilaurate (250 µM) at 60 °C for 6 hours.

*(2) Measurements using RP-HPLC-based separation of DIs and TPA.* Further, TTCE's potential to hydrolyze the following substrates was assessed and compared with LCC's potential: (i) PET films and granules, and (ii) PET's degradation intermediates (DIs), specifically bis-hydroxyethyl-terephthalate, or BHET (incubation for 12 hours, at 60 °C). For the work with PET granules, granules were either used directly with enzyme (incubation for 50 h, at 60 °C), or dissolved in hexafluoroisopropanol (HiMedia Laboratories) and then dried in an incubator-shaker at 50 °C to form a film which was washed with distilled water and phosphate buffer of pH 8.0, prior to reaction with enzyme (incubation for 4 days, at 60 °C). Activity was also assessed using commercially-sourced PET films (Goodfellow, Product code: GF25214475) of thickness 0.25 mm, which were cut into circular discs of radius 3 mm, washed with 1% SDS at 50 °C for 30 minutes, and then with de-ionized water and ethanol, using the same temperatures and time periods, before overnight air-drying of the ethanol and use in enzymatic degradation experiments (incubations for 50 h, at 60 °C). The products of degradation, namely, TPA, BHET and MHET, were quantified using a Shimadzu HPLC equipped with a photodiode array (PDA) detector and a LiChrospher® RP-18 column (5 µm

particle size, L × I.D. 25 cm × 4.6 mm), with dilution of samples in a 1: 1 ratio with acetonitrile, prior to injection of 20 µl of this dilution into the column, for separation using a mobile phase of phosphate buffer of pH 2.5 (Solvent A) and methanol (Solvent B), run conditions consisting of 3 timed stages (0-5 minutes, 25% B; 5-22 minutes, 25%-100 % B; 22-27 minutes, 25% B), and measurement of the absorbance of DIs at 240 nm, with quantification of products through reckoning of the area under each elution.

*(3)* *Scanning Electron Microscopy.* PET polymers in the form of circular films of ~6 mm diameter were examined using SEM after the enzymatic treatment. The untreated and enzyme-treated films were washed with 1% SDS, water and ethanol. Films were air dried and sputter-coated with gold of 5 nm thickness. Coated films were mounted on aluminum stubs using carbon tape, and SEM imaging was done using a Jeol Field emission Scanning Electron Microscope, with a beam accelerating voltage of 15 keV.

### Section 3. Supporting Tables

**Table S1:** Docking scores from Glide-XP-dock and binding scores from Prime-MMGBSA calculations, using **Schrödinger** software.

| **Hydrolase** | **Ligand** | **Dock Score** | **Glide emodel** | **Prime Energy** | **MMGBSA ΔG binding** |
|---|---|---|---|---|---|
| LCC | 2HE-$(MHET)_4$ | -5.06 | -74.71 | -10336.95 | -71.95 |
| TTCE | 2HE-$(MHET)_4$ | -4.87 | -80.7 | -9423.35 | -72.93 |

**Table S2:** Basic properties of TTCE, LCC, and the TTCE-LCC fusion construct.

| | Molecular weight (kDa) | Theoretical pI | Number of negatively charged residues | Number of positively charged residues | Number of tryptophan residues | UV absorbance of a 0.1% solution (1 mg/ml) at 280 nm |
|---|---|---|---|---|---|---|
| TTCE | 26.9006 | 9.29 | 26 | 29 | 1 | 0.555 |
| LCC | 29.413 | 9.57 | 15 | 22 | 4 | 1.31 |
| TTCE-LCC Fusion | 56.644 | 9.47 | 41 | 51 | 5 | 0.935 |

**Table S3:** Comparison of thermal (CD and intrinsic fluorescence), kinetic (Gdm.HCL induced unfolding measured using CD) and thermodynamic stabilities (DSC) of TTCE and LCC

| | **Apparent $T_m$ from CD** | **Red shift from intrinsic fluorescence** | **Rate of kinetic unfolding in water ($K_{u;w}$)** | **Enthalpy of unfolding using DSC (Kcal/moles)** |
|---|---|---|---|---|
| **TTCE** | No unfolding observed till 90 °C | No change observed till 90 °C | $2.6*10^{-20}$ $s^{-1}$ | ΔH = 1915.87 |
| **LCC** | 83.7 °C | 80 °C | N.D. | Two state transition $\Delta H_1$, $\Delta H_2$: 302.1, 231.9 |

### Section 4.  Supporting Figures

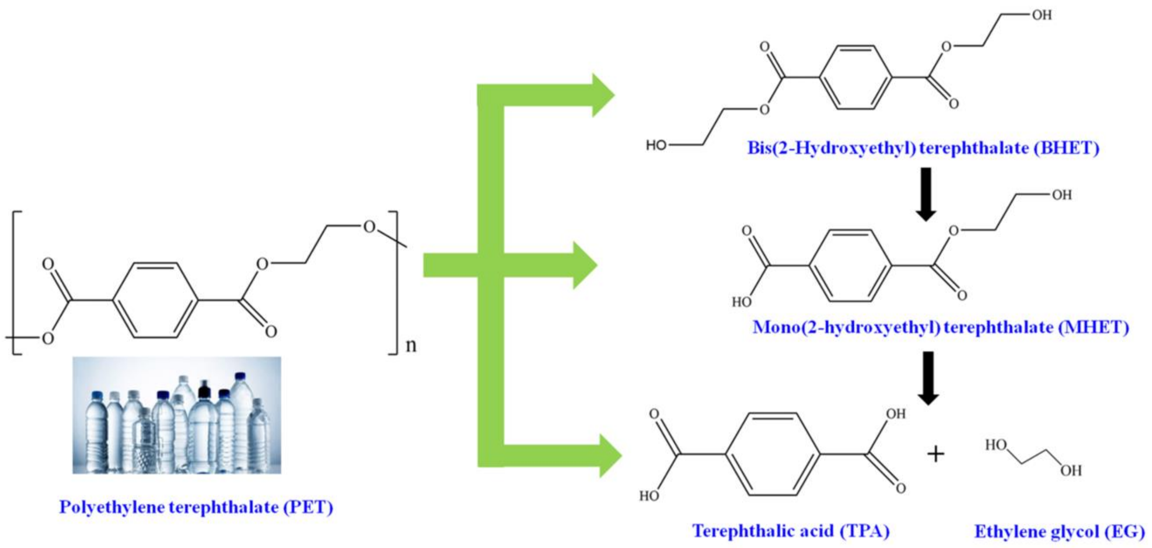


**Figure S1**: Enzymatic breakdown of polyethylene terephthalate (PET) or oligoethylene terephthalate (OET) into different degradation intermediates (DIs). In PET, the ‘n’ is large, whereas in OET (not shown) the ‘n’ is small. The ‘exolytic’ action of enzymes upon PET/OET can generate the terminal degradation product, terephthalic acid (TPA), along with ethylene glycol (EG). Exolytic action can also generate degradation intermediates such as Bis-(2-hydroxyethyl) terephthalate (BHET), and Mono-(2-hydroxyethyl) terephthalate (MHET). Multiple ‘endolytic’ actions of enzymes upon PET generate OET.

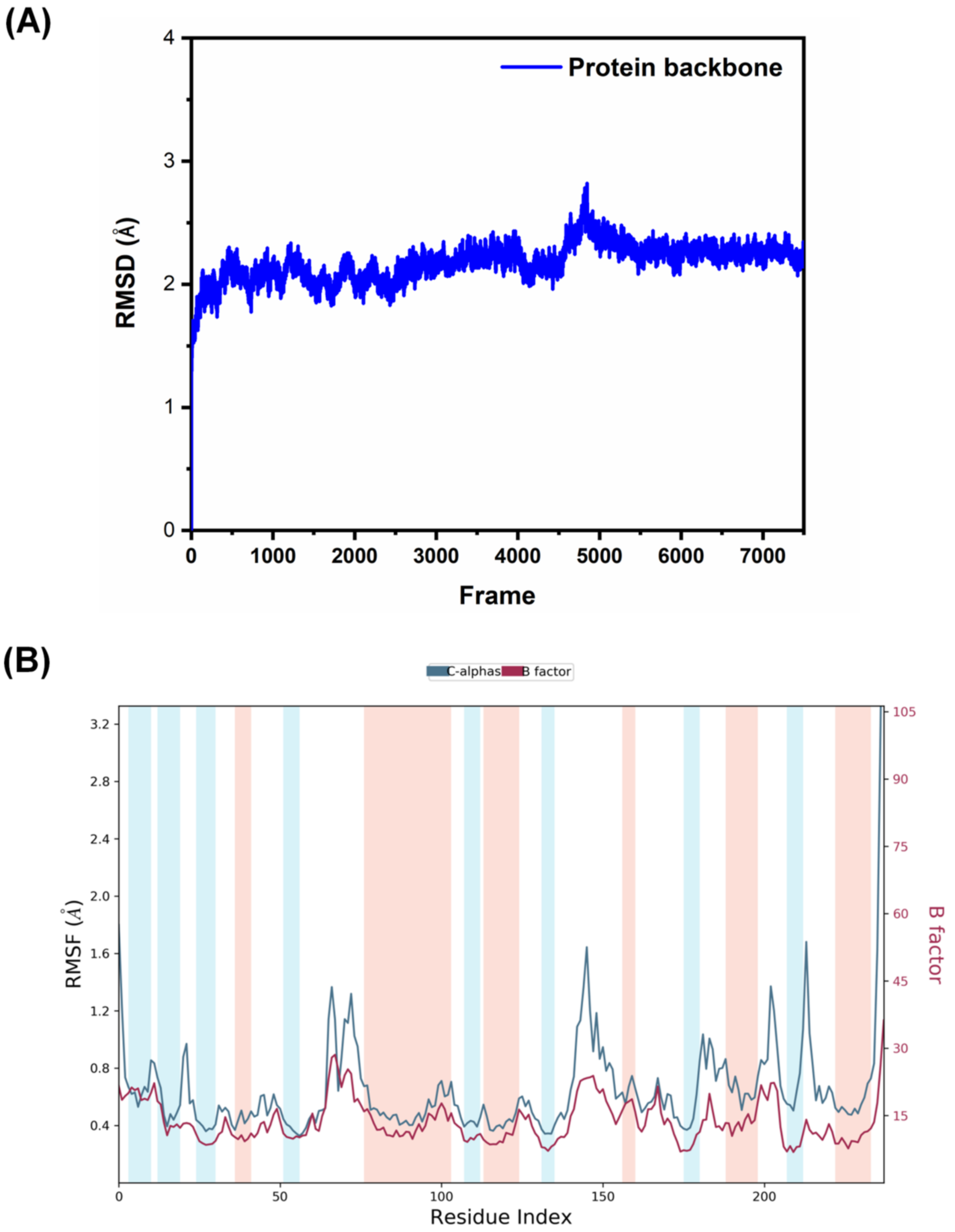


**Figure S2**: Molecular dynamics simulation of TTCE-2HE-(MHET)$_4$ for 75 ns shows (A) dynamically stable trajectory as indicated by the flattened root mean-square deviation (RMSD) in the protein structure relative to the initial structure. (B) rootmean-square fluctuations (RMSF) indicating the extent of flexibility at each residue position. The peaks denote the position of flexible residues in TTCE most likely to interact with short chain PET.

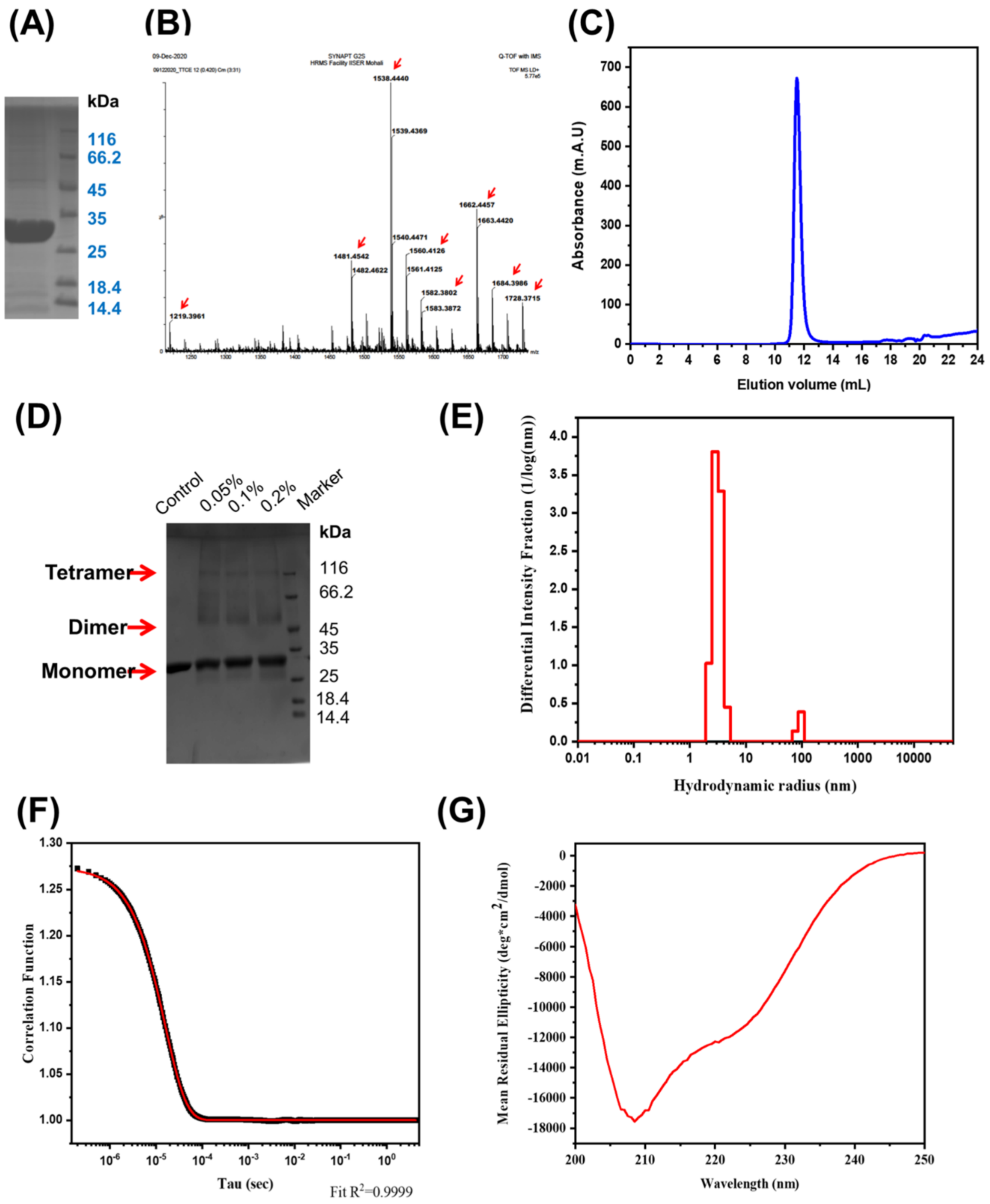


**Figure S3:** (A) SDS-PAGE lanes showing the eluted fraction obtained during purification of TTCE. (B) MALDI-Q-TOF peptide mass fingerprint of TTCE, with red arrows representing peptide masses observed that matched with masses of in silico trypsin-digested TTCE. (C) Size exclusion chromatogram of TTCE, monitoring elution of protein absorbing at 280 nm (characteristic absorption of tryptophan residues). (D) Outcome of cross-linking of TTCE by varying concentrations of glutaraldehyde. (E) Dynamic light scattering results, showing distribution of sizes of TTCE. (F) Correlation function for the light scattering results. (G) Circular Dichroism spectrum of the mixed α/β structure of TTCE.

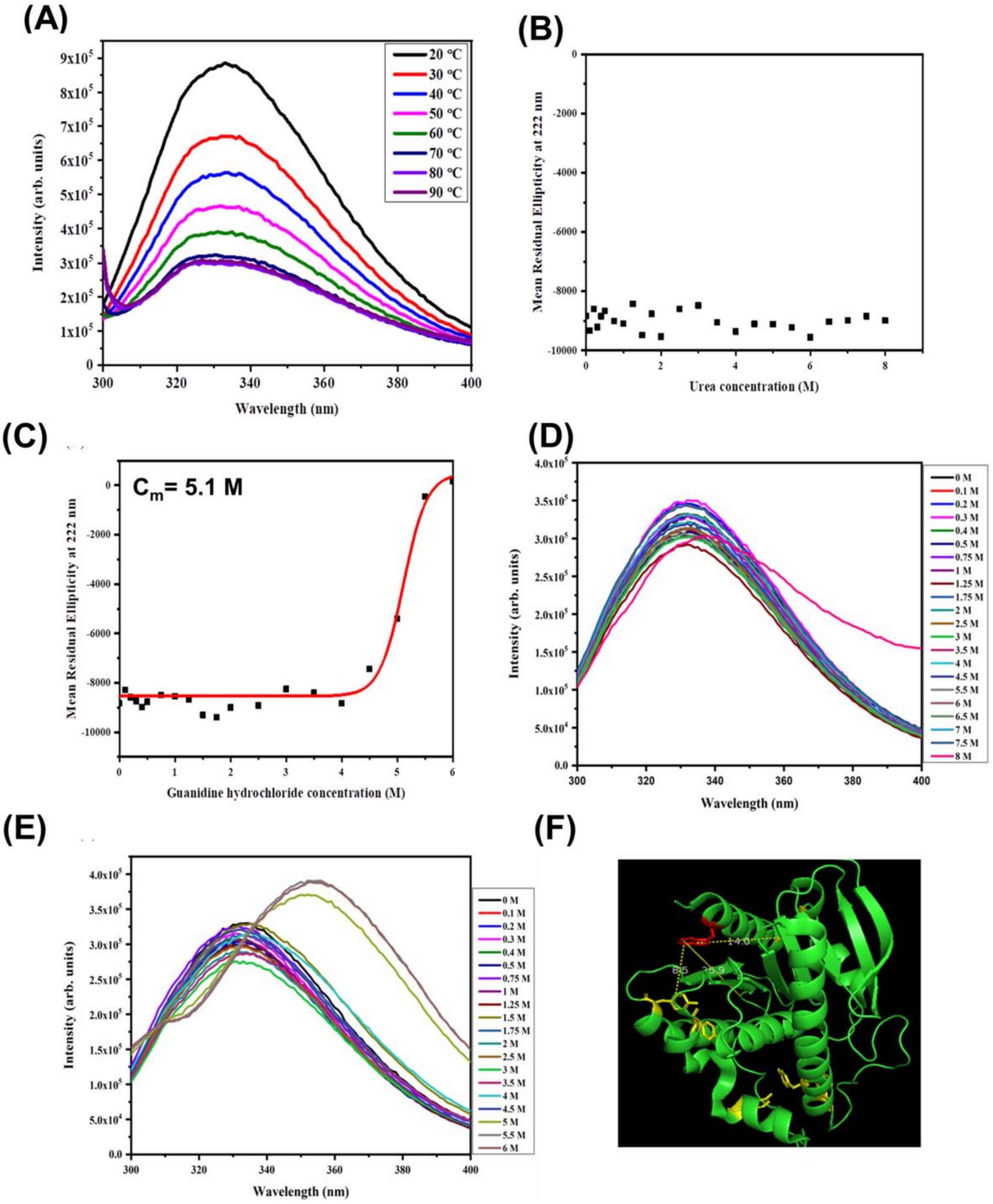


**Figure S4:** (A) Lack of changes in tertiary structure of TTCE (fluorescence spectral emission maxima) as a function of temperature. (B) Boltzmann fit showing lack of changes in secondary structure up to 8 M urea (MRE at 222 nm versus urea concentration). (C) Boltzmann fit showing lack of changes in secondary structure up to 4 M Gdm.HCl (MRE at 222 nm versus Gdm.HCl concentration). (D) Lack of changes in tertiary structure up to 7.5 M urea (fluorescence emission spectra of TTCE as a function of urea concentration). (E) Lack of changes in tertiary structure up to 4.5 M Gdm.HCl (fluorescence emission spectra of TTCE as a function of Gdm.HCl concentration). (F) Distances in Å, between the sole tryptophan in TTCE and the surrounding tyrosine residues in the folded state of TTCE, facilitating energy transfer between tyrosine (donor) and tryptophan (acceptor) in native TTCE (PDB ID: 1UFO). In TTCE subjected to unfolding, energy transfer no longer occurs, leading to (a) release of the quenching of tryptophan through FRET and intensification of tryptophan fluorescence, and (b) distinct fluorescence emissions from tyrosine at ~305 nm, and tryptophan at ~340 nm.

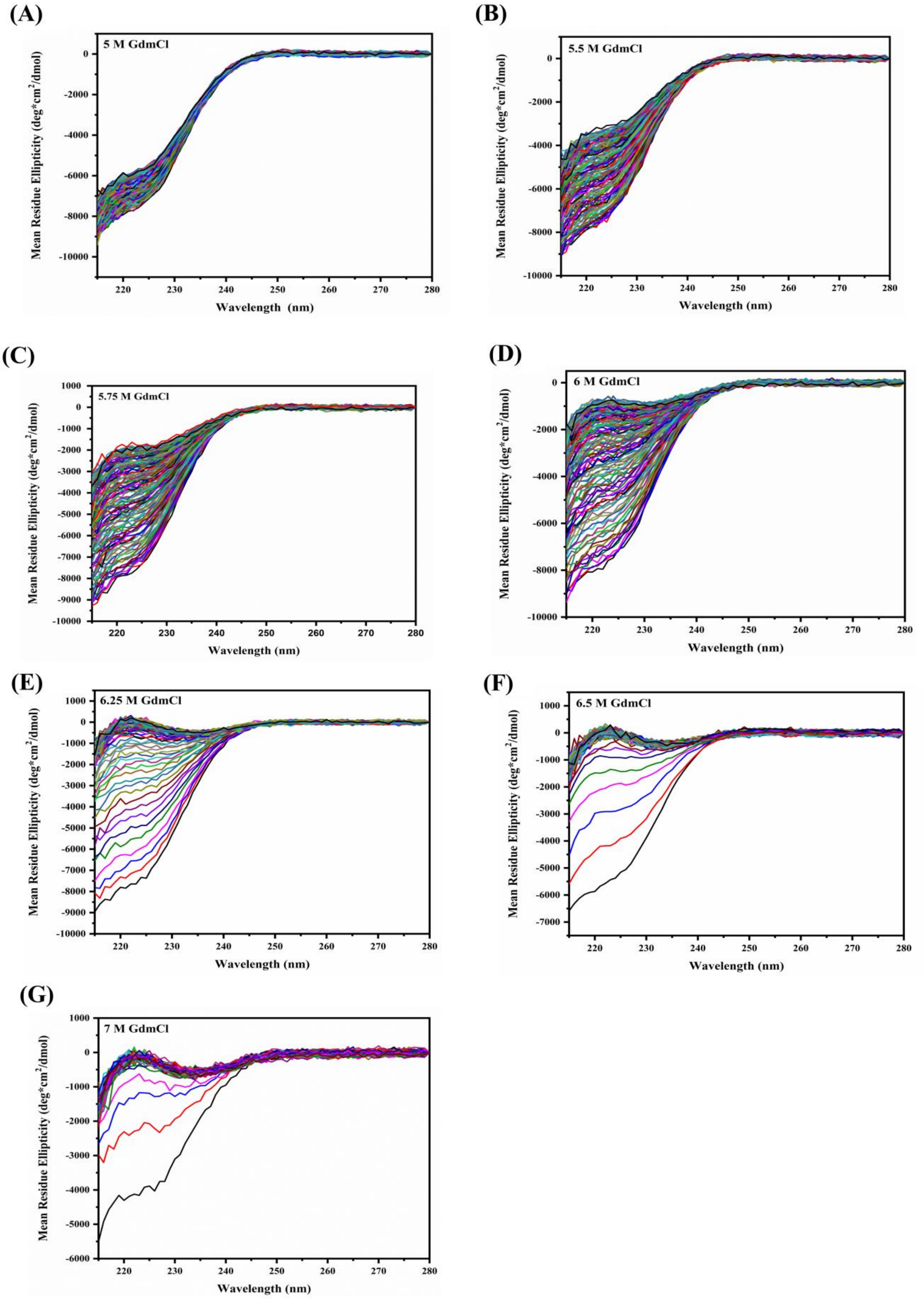


**Figure S5:** Kinetic stability studies of TTCE in the presence of Gdm.HCl (also known as Gdm.Cl). (A to G) Changes in CD spectra elicited by Gdm.HCl concentrations in the range of 5 M to 7 M, during the course of 2 hours of incubation with the denaturant, with spectra collected at 1 minute intervals.

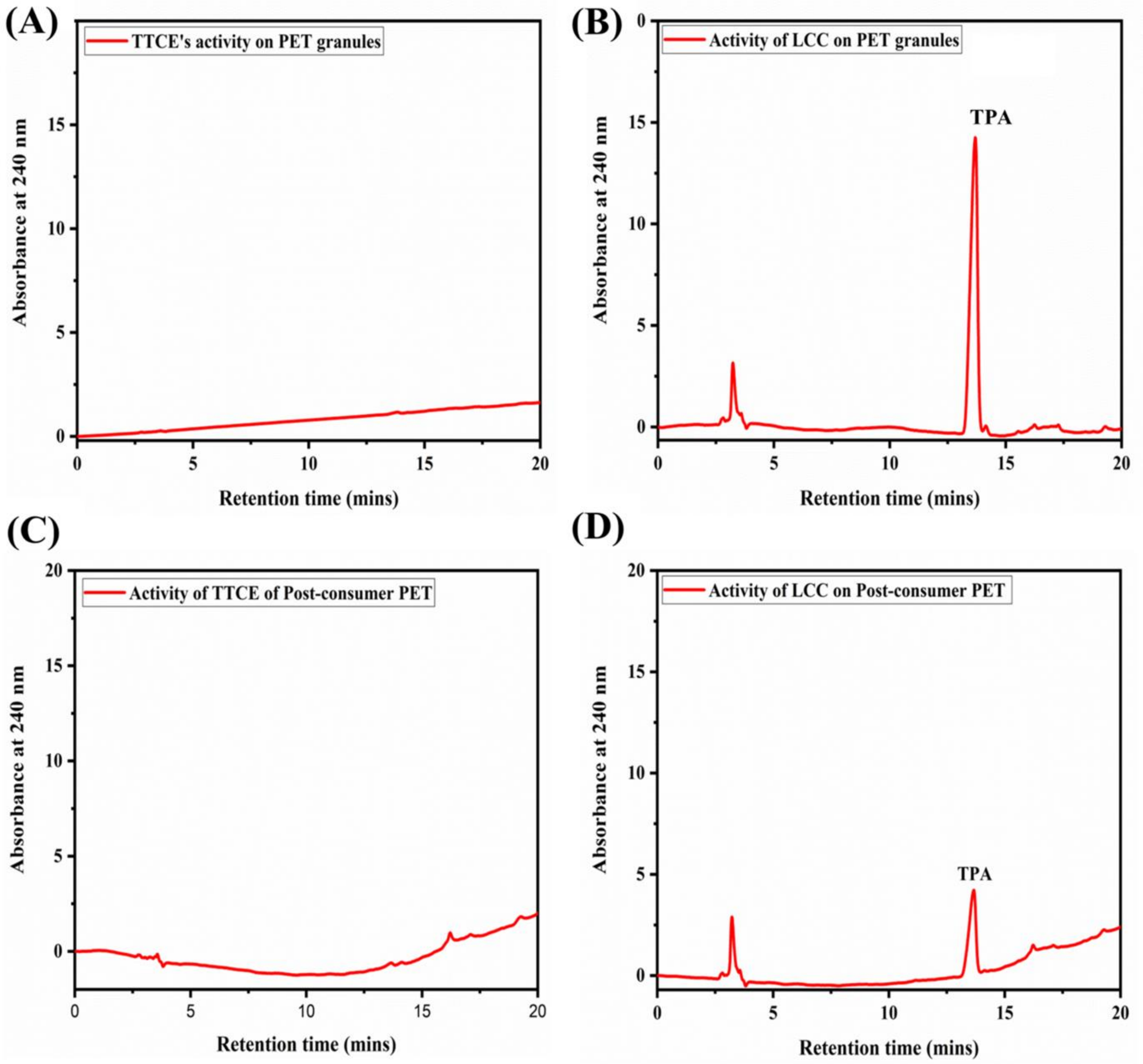


**Figure S6**: The degradation of intact PET granules by the action of (A) TTCE and (B) LCC; as well as the enzymatic hydrolysis of post-consumer PET by (C) TTCE and (D) LCC.

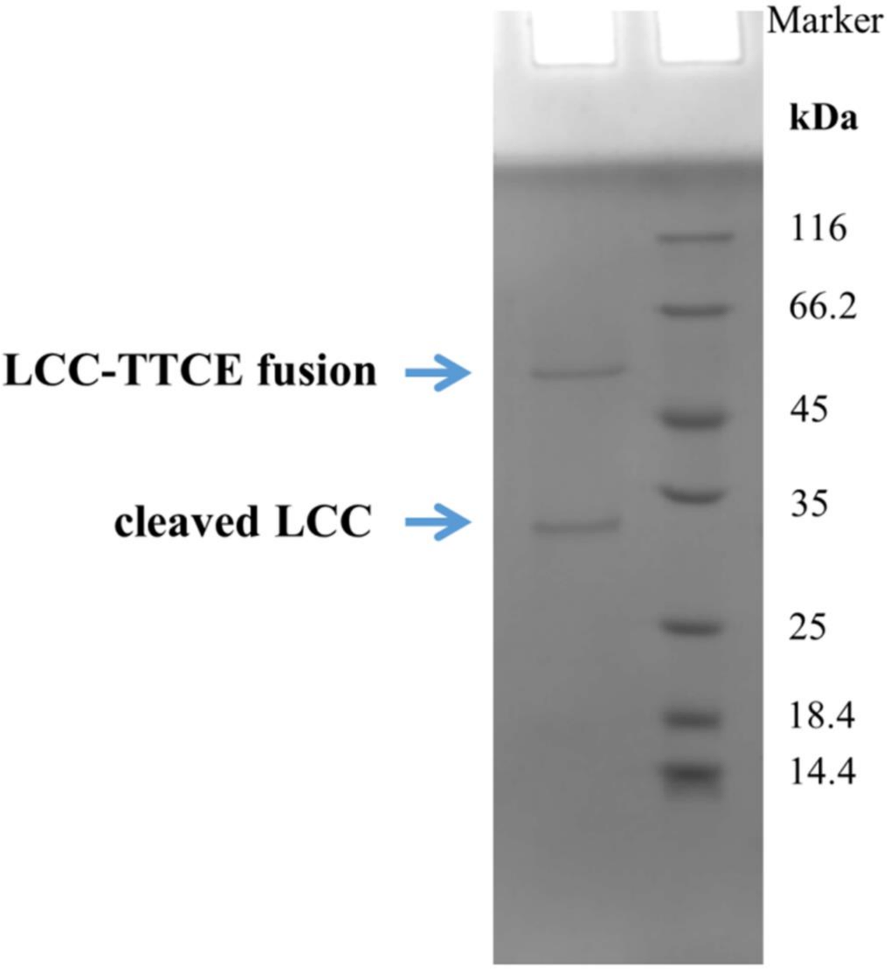


**Figure S7:** The PET-film bound fraction of the LCC-TTCE fusion, retrieved by boiling the PET films in SDS-PAGE loading buffer. The fraction shows the two populations corresponding to the intact LCC-TTCE fusion construct bound to PET, and the LCC bound to PET after the proteolytic degradation of the linker between the LCC and TTCE, suggesting the release of some molecules of TTCE in the solution.

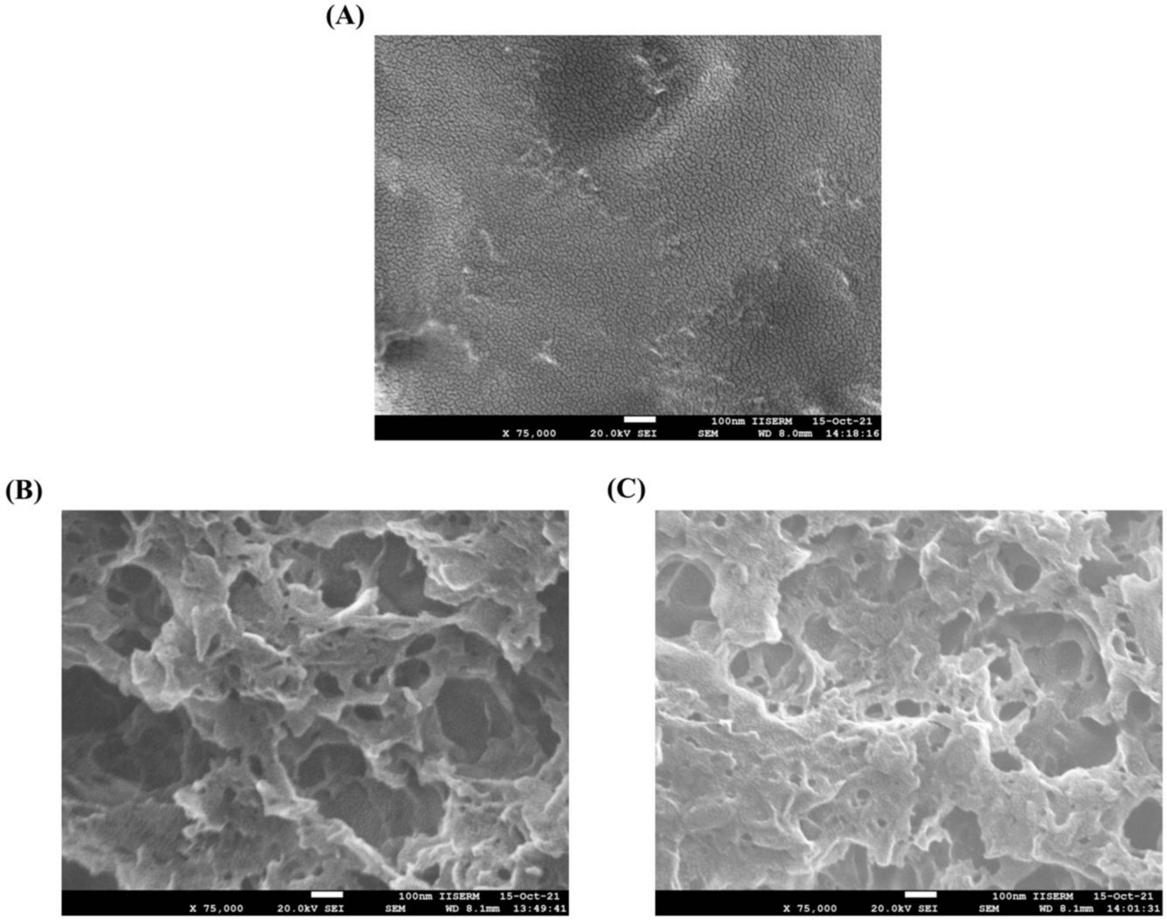


**Figure S8**: Scanning electron microscopy (SEM) images of the surfaces of PET films subjected to exposure to the LCC-TTCE fusion construct, and the LCC+TTCE enzyme cocktail. Magnified images are presented. (A) Untreated PET surface (75000X). (B) Surface treated with the TTCE+LCC enzyme cocktail (75000X). (C) Surface treated with the LCC-TTCE enzyme fusion (75000X). Greater invasion of the PET surface appears to be achieved with the LCC+TTCE enzyme cocktail and the LCC-TTCE enzyme fusion than with LCC alone.

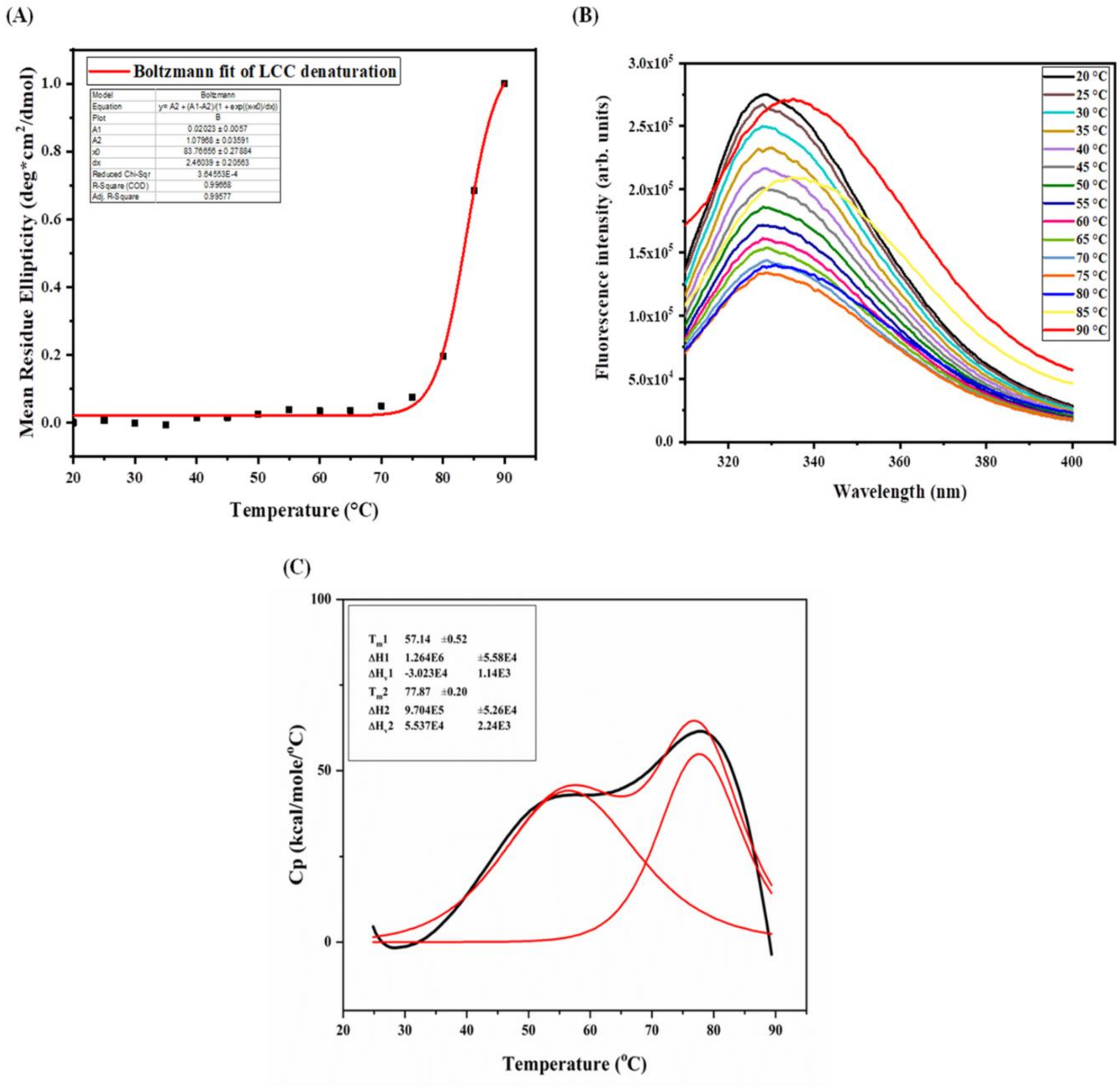


**Figure S9:** Thermal stability of LCC. (A) Lack of changes in secondary structure below 75 ˚C (assessed through measurement of Circular Dichroism spectra). (B) Lack of changes in tertiary structure below 75 ˚C (assessed through intrinsic fluorescence emission maxima). (C) Differential scanning calorimetry profile of LCC, showing an enthalpic transition with a $T_m$ of ~ 77.87 ˚C, with significant enthalpic changes also seen at ~ 57 ˚C (however, without effects upon secondary or tertiary structure, as suggested by panels A and B of this figure).
